# Predicting microbial transcriptome using annotated genome sequence

**DOI:** 10.1101/2024.12.30.630741

**Authors:** Guohao Fu, Yujing Yan, Ziyue Zhao, Ye Chen, Bin Shao

## Abstract

We present TXpredict, a transformer-based framework for predicting microbial transcriptomes using annotated genome sequences. By leveraging information learned from a large protein language model, TXpredict achieves an average Spearman correlation of 0.53 and 0.62 in predicting gene expression for new bacterial and fungal genomes. We further extend this framework to predict transcriptomes for 2, 685 additional microbial genomes spanning 1, 744 genera, 82% of which remain uncharacterized at the transcriptional level. Our analysis highlights conserved and divergent transcriptional programs across understudied genera, providing a powerful resource for uncovering microbial adaptation strategies and metabolic potential across the tree of life.

**GRAPHICAL ABSTRACT:** 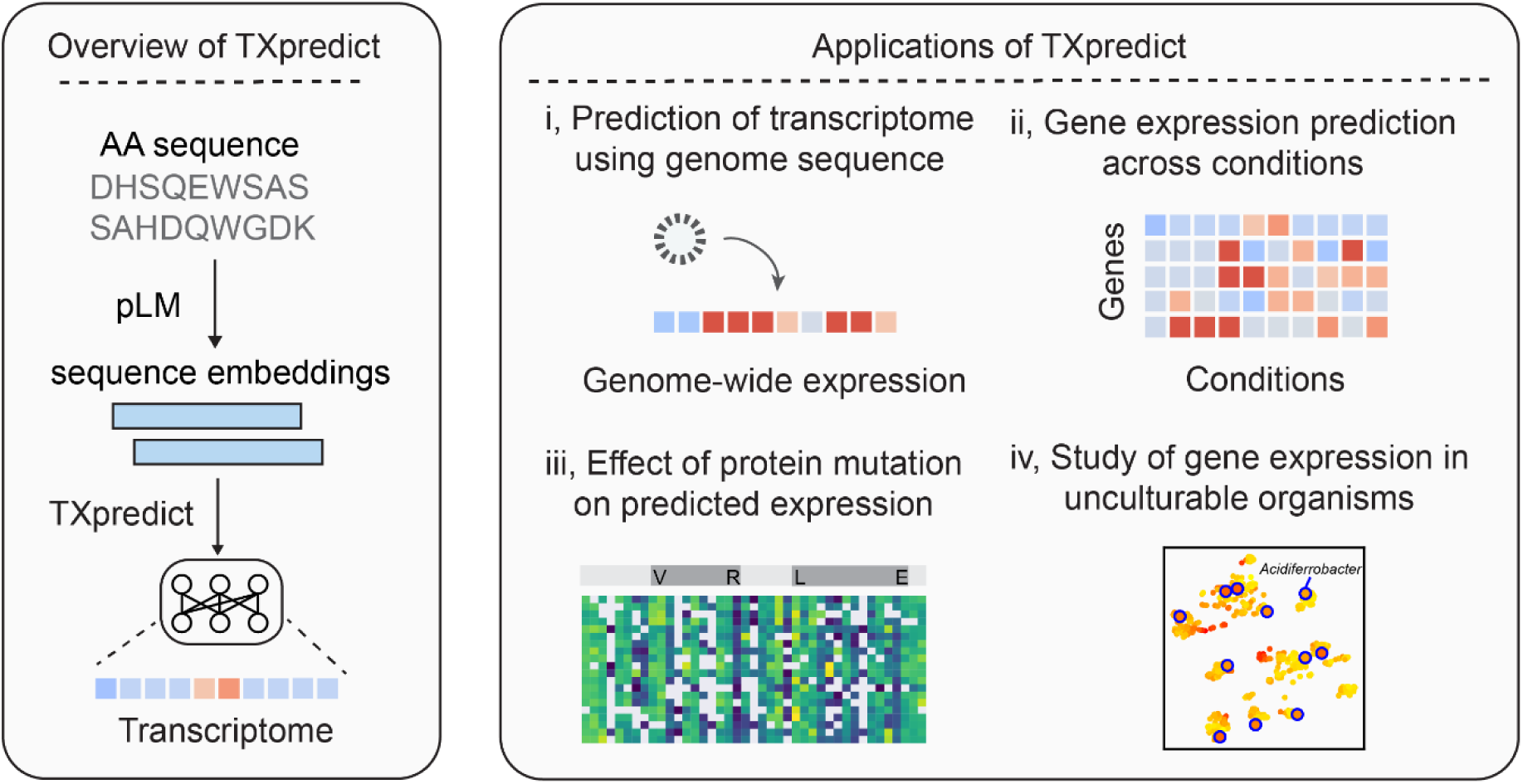

## INTRODUCTION

For a genome-sequenced microbe, transcriptome profiling provides critical insights into its adaptation, survival, and pathogenicity, and bridges the gap between raw genomic data and functional biology(1–6)(. Despite its importance, only a small fraction of sequenced microbes has been characterized at the transcriptional level. A major challenge is that most microbes remain uncultivated due to the limited knowledge of their growth requirements or their inhabitation of extreme environments(7). Recent advances in predicting physicochemical requirements have begun to address these challenges, representing an important first step(8–10). Moreover, even when cultivation is successful, RNA-seq protocols often require organism-specific optimizations, such as cell lysis and rRNA depletion, which can involve extensive trial and error for novel organisms. These technical issues hinder large-scale, cross-species studies of microbial gene expression across the tree of life.

Current machine learning approaches for predicting gene expression have primarily focused on analyzing untranslated regions (UTRs)(11–16) or large genomic windows surrounding the gene of interest (17, 18). While these methods show promise, their application to non-model organisms remains limited, as trained models often fail to generalize across species or even within the same species(19, 20). Additionally, previous studies have demonstrated evolutionary constraints on protein synthesis, such as conserved enzyme stoichiometry in metabolic pathways(21–23). Yet these constraints have not been translated into predictive models that are broadly applicable to diverse microbial species.

Here we introduce TXpredict, a novel framework for predicting microbial transcriptomes using only annotated genome sequences. Our approach uses the protein-coding sequences derived from the genome as the model’s input, and it leverages a protein language model, ESM2(24), to extract predictive features. Using transcriptome data from 22 bacterial, 9 archaeal and 9 fungal species obtained from the NCBI GEO database(25) as the ground-truth target, we trained a transformer-based model that achieves a Spearman correlation up to 0.70 in leave-one-genome-out evaluations. We validated the model’s performance in the non-model organism *Pseudomonas sp*. WBC-3 by conducting a series of RNA-seq experiments. By leveraging large-scale measurements from Deep Mutational Scanning (DMS) assays, we found that TXpredict’s predictions of mutant sequence expression correlate with experimentally measured protein stability for specific proteins. Lastly, we applied the trained TXpredict models to predict transcriptomes for 2, 685 additional genomes, the majority of them have never been transcriptionally characterized. This resulted in the creation of TXpredictDB, a database containing expression profiles for 9.8 million genes. The models developed in this study are publicly available: https://github.com/lingxusb/TXpredict

## MATERIAL AND METHODS

### Training dataset

We collected RNA-seq data from the NCBI GEO database (https://www.ncbi.nlm.nih.gov/geo/). For each microbe species, we selected the dataset with the highest number of samples and used either raw count data or RPKM (Reads Per Kilobase of transcript per Million mapped reads) values when available. For raw count data, we calculated the RPKM for each gene, followed by log (1 + p) and z-score normalization. In total, we gathered data from 22 bacterial species (2, 649 samples), 9 archaeal species (169 samples) and 9 fungal species (2, 078 samples, Supplementary Table 1). For each species, we constructed a standard gene expression profile by averaging the normalized expression values across all available samples.

To model gene expression across conditions, we also collected RNA-seq data from iModulonDB (https://imodulondb.org/index.html). We downloaded the processed datasets for each microbial species, resulting in a dataset that includes 13 bacterial species and a total of 4, 619 conditions. For each microbial species, we only kept the genes with mean normalized gene expressions greater than −3 and a standard deviation less than 1 across conditions.

Protein embeddings for all annotated genes were calculated using the ESM2-650M model (*esm.pretrained.esm2_t33_650M_UR50D()*), following the instructions provided at https://github.com/facebookresearch/esm. Proteins longer than 1, 500 amino acids (AAs) were excluded from this analysis. In addition to the protein embeddings, we also included basic protein sequence statistics in the model input: the normalized protein length (calculated by dividing each protein length by the longest protein length within each organism) and the amino acid composition (the proportion of each of the 20 amino acids). The final input dimension of the model is 1301.

### Processing of Nanopore sequencing datasets

We downloaded and processed nanopore sequencing datasets for three bacteria strains: *Pseudomonas aeruginosa* UCBPP-PA14 (PMID: 34780302)(26), *Haloferax volcanii* DS2 (PMID: 37192814)(27), and *Escherichia coli* str. K-12 substr. MG1655 (PMID: 39011882; 39453698; 34906997)(28–30). The analysis was performed with Miniconda3, minimap2 (v2.24) for read alignment, samtools (v1.16) for processing alignment files, and featureCounts (from the Subread package v2.0.6) for read counting. Sequence Read Archive (SRA) files for each sample (e.g., SRR27228856) were downloaded from the NCBI SRA database using the prefetch command. SRA files were converted to FASTQ format using fasterq-dump. Sequencing reads were aligned to the reference genome using minimap2 with the -ax map-ont parameter, optimized for Oxford Nanopore long-read data. The resulting SAM files were converted to BAM format, sorted, and indexed using samtools. Alignment quality was assessed using samtools flagstat to generate mapping statistics. Read counts for each gene were generated from the sorted BAM files using featureCounts with the gene annotation file (GTF) as a reference. The resulting count tables were then processed using custom awk scripts to map gene identifiers to official gene names, producing a final gene expression matrix for each sample. To account for biases in gene length and sequencing depth, we normalized the gene counts as previously described. Genes not detected in any replicate were excluded from further analysis.

### Transcriptome prediction model

We developed a Transformer-based model to predict gene expression levels from sequence features. The transformer encoder utilizes a multi-head self-attention mechanism (num_heads=4) with a single encoder layer. The encoder processes input features through a linear transformation into a hidden dimension of 128, followed by positional encoding. We implemented sinusoidal positional encoding to capture sequential information. The output of the transformer layer was then fed into two fully connected layers (dimension 512 and 128) with ReLU activations. The final linear layer produces a single output value representing the predicted gene expression level.

### Model training and evaluation

We performed cross-species evaluations for bacteria, archaea and fungi separately to account for potential domain-specific expression patterns. For each domain, we employed a cross-validation strategy where models were trained on data from all but one microbial species and tested on the held-out species. This approach was repeated for each species in our dataset to assess our model’s generalization capability. The models were trained using the Adam optimizer with a learning rate of 0.0002 for 5 epoch with a batch size of 32. To ensure robust evaluation, we performed 5 independent training runs with different random seeds (42, 123, 456, 789, 1011). The mean squared error (MSE) loss function was used for optimization. Model performance was evaluated using both Spearman and Pearson correlation coefficients between predicted and measured expression levels.

New bacterial, archaeal and fungal genomes were downloaded from NCBI Genbank (https://ftp.ncbi.nlm.nih.gov/genomes/genbank/). To maximize genus-level representation for transcriptome prediction, we selected 2, 169 bacterial, 423 archaeal, and 93 fungal genomes spanning a total of 1, 744 genera. Gene expression for these genomes was predicted using TXpredict models, forming the basis of the TXpredictDB. To determine whether a genus has been transcriptionally characterized, we searched the GEO database using the following query: “*genus_name [Organism] AND ("Expression profiling by array"[Filter])”*. We annotated taxonomy of these genomes by querying the GTDB database(31). For model training and inference, we used 3090 Ti GPU (24GB) and the software packages PyTorch (version 2.1.1) and transformer (version 4.28.1).

### Conditional gene expression model

The conditional gene expression model shares the same transformer encoder architecture with the transcriptome prediction model, and was trained separately for each species. The UTR branch processes one-hot encoded sequences from −100 to +100 bp relative to the transcription start site. The transformer layer has 2 attention heads and a feedforward dimension of 128. The protein embedding branch utilizes 8 attention heads with a hidden dimension of 256. The outputs from both branches are concatenated together and processed through a fully connected layer (dimension: 4096) with ReLU activation. The output dimension of the model is the number of conditions for each bacterial species.

For each bacterial species, we trained the model for 10 epochs using the Adam optimizer with a learning rate of 0.0002 and a batch size of 32. Model performance was evaluated using 5-fold cross-validation. Pearson correlation coefficients were calculated between predicted and measured expression levels across all conditions for genes in the test set.

### Analysis of protein sequences

To visualize the high-dimensional embeddings in a two-dimensional space, we implemented t-SNE (t-Distributed Stochastic Neighbor Embedding) using the opentsne package. We initialized the embedding using PCA and constructed a uniform affinity matrix with 10 nearest neighbors. The optimization process consisted of three phases: an early exaggeration phase of 125 iterations with an exaggeration factor of 12 and momentum of 0.5, followed by a gradual exaggeration annealing phase where the exaggeration factor was linearly decreased from 12 to 1 over 125 iterations with momentum of 0.8. The final optimization phase ran for 2000 iterations without exaggeration. Function prediction for these proteins was carried out using the default deepFRI model(32), as available on GitHub (https://github.com/flatironinstitute/DeepFRI). We used a score cutoff of 0.5 which was reported to be significant in the original publication. To cluster the protein sequences in TXpredictDB, we used the MMseqs2(33) package with default settings and a minimum sequence identity threshold of 0.9.

### Calculation of input gradient

To investigate the contribution of individual nucleotides in the UTR sequence to gene expression prediction across conditions, we implemented a gradient-based analysis. Using a sequence-only model variant where the protein embedding branch was masked, we calculated the gradient of the model output with respect to the input DNA sequence. For each target gene and condition, we performed a forward pass with the one-hot encoded UTR sequence while setting all other inputs to zero. We then computed the gradients through backpropagation and multiplied them element-wise with the input sequence (Gradient × Input method)(17). The absolute values of these products were summed across the four nucleotide channels (A, T, C, G) to obtain single-base resolution importance scores. For visualization purposes, we aggregated these scores into 5-bp bins by summing the scores within each bin.

### RNA-seq experiment of the *Pseudomonas* sp. WBC-3 strain

For the RNA-seq experiment, *Pseudomonas* sp. WBC-3 was cultured under three different conditions: PNP1 (minimal medium with 0.01% glycerol), PNP2 (minimal medium with 0.04% PNP), and PNP3 (minimal medium with 0.01% glycerol and 0.04% PNP). Each condition was prepared by diluting a 2× stock of yeast extract minimal medium supplemented with either 0.02% glycerol, 0.08% PNP, or both, resulting in the desired final concentrations. WBC-3 strain was first grown overnight in LB medium, then diluted 1:100 into 4 mL of each experimental medium and incubated overnight at 30°C with shaking at 200 rpm. Cultures were harvested at OD600 between 0.5 and 1.0 by centrifugation for 2 minutes, ensuring that the total collected biomass did not exceed 1×10⁹ cells. Total RNA was extracted using the RNAprep Pure Bacteria Kit (TIANGEN, DP430), followed by DNase I treatment (TIANGEN, RT411) to remove genomic DNA contamination. RNA quality and concentration were confirmed to meet sequencing requirements. Raw sequencing reads were aligned and processed using standard RNA-seq pipelines. Read counts were quantified using StringTie with the following command: stringtie -p 8 -G GCF_035758525.1_ASM3575852v1_genomic.gtf -l sample_id -o sample.gtf sample.bam, using the *Pseudomonas* sp. WBC-3 reference genome (GCF_035758525.1).

## RESULTS

### Prediction of transcriptome based on the genome sequence

We developed two distinct models that address different aspects of gene expression prediction. The transcriptome model predicts each gene’s mean expression across diverse experimental conditions, providing a baseline expression profile. The condition-dependent model incorporates experimental context to predict how individual genes respond to different conditions. For the transcriptome model, we first collected microbial RNA-seq datasets from the NCBI GEO datasets and selected the dataset with the highest number of samples for each microbial species. Read counts were normalized with the same pipeline (Methods). Taken together, our dataset includes 22 bacterial, 9 archaeal and 9 fungal species and a total of 24.8M gene expression measurements (Supplementary Table 1). For each microbial species, gene expression was averaged across all samples to construct a standard expression profile. We found that this standard expression profile is robust to differences in condition coverage, since mean expression profiles from independent GEO datasets of the same species are strongly correlated (Supplementary Fig. 1). Protein embeddings for each gene were computed using the ESM2-650M model and combined with basic protein statistics to serve as the input features for TXpredict (Fig. 1a, Methods). For model evaluation, we employed a leave-one-genome-out cross-validation strategy. For bacterial species, the model was trained on genes from 21 species and tested on the held-out species (Fig. 1b). We followed the same approach for archaeal and fungal species.

**Figure 1.**
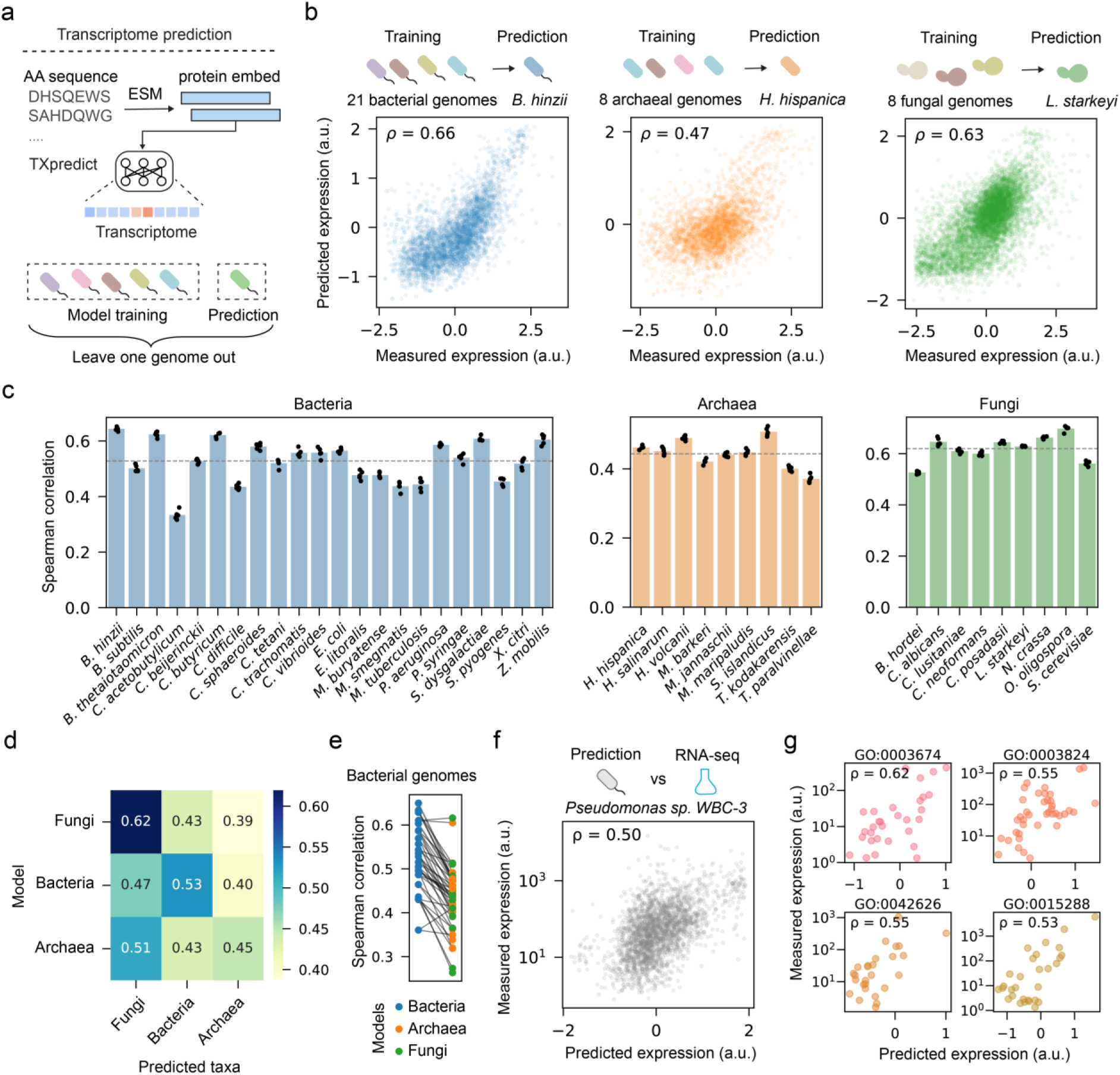
Prediction of transcriptome based on the genome sequence. **a)** Schematic of TXpredict for transcriptome prediction. **b)** Scatter plots showing predicted and measured gene expression in leave-one-genome-out evaluations for one species from bacteria, archaea, and fungi. Spearman correlation coefficients are reported in the top-left corner of each plot. **c)** Spearman correlation coefficients between predicted and measured gene expression in leave-one-genome-out evaluations. TXpredict models were separately trained for bacterial, archaeal, and fungal species. Results from five rounds of model training and testing are reported (n = 5). Dashed lines indicate the mean correlations for each domain. **d)** Cross-domain prediction performance of TXpredict models. Models were trained on all species from one domain and tested on species from the other domains. Mean Spearman correlations across all test species are shown. Diagonal values represent leave-one-genome-out performance within each domain. **e)** Predictions of archaeal and fungal models on all bacterial species. Blue dots indicate leave-one-genome-out evaluations for bacteria. Predictions for the same species were connected. **f)** Model predictions versus measured gene expression in new RNA-seq experiment for *Pseudomonas sp.* WBC-3. Each dot represents one functional gene and the Spearman correlation is shown. **g)** Predicted and measured gene expression for genes with different GO annotations in *Pseudomonas sp.* WBC-3.

We found our model achieved a mean Spearman correlation coefficient of 0.53 for the bacterial species (Fig. 1c), and performed well for non-model organisms, with correlation coefficients of 0.64, 0.62 and 0.62 for *B. hinzii*, *B. thetaiotaomicron*, and *C. beijerinckii*. It is worth noting that transcriptome predictions for these bacteria were based solely on the annotated genome sequences. The model’s performance in predicting gene expression variation decreased modestly as the variation calculations depend on the diversity of experimental conditions (Supplementary Fig. 2). Despite having a limited training set for fungal species, our TXpredict model obtained a mean correlation of 0.62 and reached a Spearman correlation coefficient up to 0.70 for non-model organisms such as *O. oligospora* (Fig. 1c). For archaea, TXpredict showed a mean Spearman correlation of 0.44 (Fig. 1c) and demonstrated broad applicability to extremophiles from distinct natural niches (ranging from the Dead Sea to volcanic springs), further highlighting the robustness of our method.

The model performance for the test genome has a weak correlation with its evolutionary distance to the training genomes (Supplementary Fig. 3), while adding UTR features into model input didn’t significantly improve model performance (Supplementary Fig. 4). Even though our model is trained on proteins shorter than 1500 amino acids, it interpolated well to proteins longer than 1500 AA, especially for yeast proteins (Supplementary Fig. 5). Lastly, we evaluated model performance within functional categories. Expression levels of test genes sharing the same GO annotation can still vary considerably. TXpredict captures this fine-scale variation, with within-group correlations only slightly lower than that for all genes. (Supplementary Fig. 6).

In our initial evaluation, TXpredict performed poorly on the archaeon *N. maritimus*, with a Spearman correlation of 0.23 in the leave-one-genome-out evaluation (Supplementary Fig. 7). Upon closer examination of this dataset, we found that fewer than 40% of the reads were successfully mapped to reference genome, demonstrating the potential of our model to detect potential problems in RNA-seq data.

To evaluate whether TXpredict merely memorizes training data or learns generalizable patterns, we examined whether protein sequence similarity affects model performance. For bacterium *B. hinzii*, we used BLASTP(34) to compute sequence similarity between its proteins and those in the other bacterial strains. The model was trained on other species and tested on rare proteins from *B. hinzii* that were not present in the training data, showing a correlation of 0.46 for them (Supplementary Fig. 8). While this is lower than the correlation for all proteins (Fig. 1c), it suggests that the model captures expression-related features beyond sequence similarity. We also benchmarked the performance of our model with a BLASTP-based nearest-neighbor baseline. For each protein in the held-out test genome, we used BLASTP to identify the most similar sequence in the training set and transferred its expression value as the prediction. BLASTP identified close homologs (with significant matches) for >95% of test proteins. For these proteins, TXpredict achieves higher Spearman correlations than the nearest-neighbor baseline in 21 out of 22 bacterial genomes, 6 out of 9 archaeal genomes, and all fungal genomes. The mean Spearman correlation values for TXpredict are 0.53 (bacteria), 0.44 (archaea) and 0.62 (fungi), compared to the nearest-neighbor baseline values of 0.38, 0.37 and 0.39, respectively (Supplementary Fig. 9). For the small fraction of sequences lacking BLASTP matches to the training set, TXpredict maintains robust performance comparable to performance on sequences with homologs, suggesting that TXpredict learns meaningful biological relationships between sequence features and expression levels.

We next explored TXpredict’s cross-taxa performance by training separate models on bacterial, archaeal, or fungal species and then testing them on other domains (Fig. 1d). Our model still showed moderate performance on cross-taxa predictions, indicating that they encoded general principles of gene expression regulation. Interestingly, the archaeal model achieved a higher correlation for eukaryotic species than on archaea. For cross-domain evaluation, the archaeal model was trained on the complete archaeal dataset before testing on other genomes. In contrast, for the archaeal leave-one-genome-out evaluation, the small number of total conditions means that removing a genome often removes unique experimental conditions, forcing the model to predict expression under unseen contexts and reducing generalization. At the species level, models trained on other taxa generally performed worse than those trained on the same taxon (Fig. 1e and Supplementary Fig. 10). To validate the prediction of the TXpredict model, we conducted new RNA-seq experiments on *Pseudomonas sp*. WBC-3 and compared the predicted expression levels with the measured values. The Spearman correlation between measured and predicted expression was 0.40 for all proteins and 0.50 for proteins with GO annotations (Fig. 1f). Notably, our model showed high correlations (0.55) for certain GO categories including catalytic activity, and also for operons like the flagellar genes (Fig. 1g and Supplementary Fig. 11).

### Robustness of the transcriptome prediction model

To evaluate the robustness of our model training process, we examined its performance against variations in the number of training species, the number of samples per species, and the sequencing platform. First, we conducted a systematic scaling analysis using bacterial datasets. For each held-out genome, we trained TXpredict on random subsets of 5, 10, and all 21 available species. The results show a modest increase in mean Spearman correlation with training species: 0.48 for 5 species, 0.50 for 10 species, and 0.53 when using all 21 training species (Supplementary Fig. 12). This scaling trend indicates that expanding the number of training species consistently improves cross-species prediction accuracy, although the magnitude of improvement diminishes with each additional species. Second, we examined whether the number of samples per species impacts performance by conducting a subsampling analysis. We randomly selected subsets of samples for each species, recalculated mean expression values, and performed leave-one-genome-out cross-validation. The results show that model performance remains stable across different sample sizes (Supplementary Fig. 13). Last, we processed and analyzed nanopore datasets including *E. coli, P. aeruginosa, and H. volcanii*, and compared these data with our Illumina-based datasets and model predictions. We observed a strong correlation in mean expression for the three strains across the two platforms (Supplementary Fig. 14). Our TXpredict model trained on Illumina data performs well for *E. coli* but poorer on *H. volcanii*, likely due to limited experimental diversity in the available nanopore dataset (n = 2).

To test whether a codon-aware approach could further improve predictions, we conducted a comparison between protein-based and RNA-based models. We used CodonTransformer(35), a transformer model that encodes DNA sequences while preserving codon information. Specifically, we used CodonTransformer to generate embeddings from coding DNA sequences of bacterial genes, and we then benchmarked this transcriptome model using the same cross-validation tests as the ESM-2 model. We found protein-based models using ESM-2 embeddings consistently outperform codon-based models (Supplementary Fig. 15).

### Modeling context-dependent gene expression

We next extended our framework to model condition-dependent gene expression (Fig, 2). We collected RNA-seq profiles from 13 bacterial strains across 4.6k unique conditions from iModulonDB, resulting in a total of 12.0M measurements (Supplementary Table 2). The model integrated one-hot encoded 5’UTR sequences with protein embeddings as input features. For each gene, we calculated the correlation between measured and predicted expression values across all conditions. Our TXpredict model achieved an average correlation coefficient of 0.52 across all bacterial strains. For *E. coli*, the distributions of Pearson’s and Spearman’s correlation coefficients show similar patterns, with 41% and 38% of genes having correlation coefficients above 0.5. The poor model performance for *P. syringae* could be due to lack of condition diversity in this dataset (Supplementary Fig. 16). Our subsampling analysis reveals a modest increase in model performance when more conditions are added (Supplementary Fig. 17). Moreover, we compared our model’s performance with a baseline nearest-neighbor model. For this baseline, we identified each test gene’s most similar sequence in the training set via BLASTP and used that gene’s expression profile as the prediction. This baseline shows an average Spearman correlation of 0.20 between predicted and measured expression across all genomes, which is much lower than that of TXpredict (Supplementary Fig. 18).

To further interpret the model’s predictions, we calculated input gradients(17) to identify sequence motifs contributing to gene expression variation across conditions (Fig. 2d and e). In *E. coli*, we observed a peak in the input × gradient overlap with the −10 box of the sigmaS-dependent promoter of the *yadV* gene(36). Also input gradients increased after the transcriptional start site, suggesting potential contributions from transcribed sequences. In a non-model organism *P. putida*, we identified a peak overlapping with a Zur binding motif for the *PP_0120* gene, consistent with its role as a Zur iModulon gene(1)(1). These findings illustrate our model’s ability to capture condition-specific gene expression changes in an interpretable manner.

**Figure 2.**
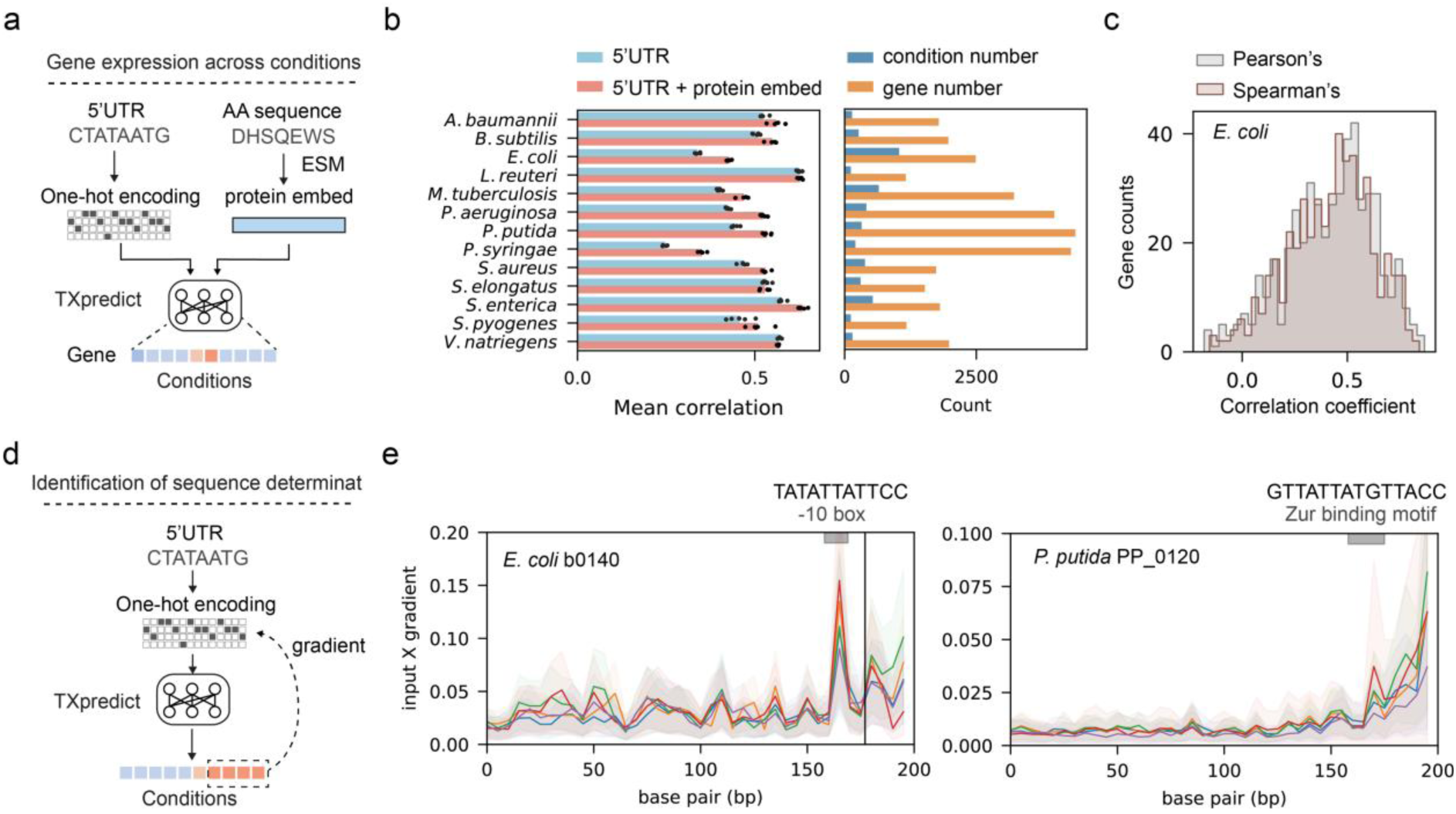
Prediction of context-dependent gene expression. **a)** Schematic of TXpredict for condition-specific gene expression prediction. **b)** Mean correlations for models trained separately on different bacterial species. Correlations from 5-fold cross-validation tests are shown on the left (n = 5). Gene and condition counts for each species are shown on the right. **c)** Distribution of Spearman and Pearson correlation coefficients for gene expression across conditions in *E. coli* (n = 498). **d)** Calculation of input gradients for identifying sequence determinants of condition-dependent gene expression. **e)** Two examples of input × gradients and their correlations with known sequence motifs.

### Relationship between protein expression and mutation effects

Deep Mutational Scanning (DMS) assays have been widely used to map protein fitness landscapes by large-scale measurement of phenotypic effects of protein mutants(37, 38). We leveraged the comprehensive benchmarks provided by ProteinGym(39) to evaluate whether predicted expression changes for protein variants correlate with experimentally measured functional scores from DMS assays (Fig. 3a). We selected yeast protein datasets, as our fungal model achieved the highest accuracy in leave-one-genome-out evaluations. We found that the correlation between protein expression and mutational effects is generally weak and protein-specific. For specific proteins, such as THO1 and MYO3, there are strong correlations between predicted expression and experimentally measured mutant effects (Fig. 3b and c). We then compared TXpredict predictions against zero-shot predictions from several machine learning-based protein models(40–47) (Fig. 3d). TXpredict’s performance was only moderately lower than two inverse-folding models (ESM-IF1(45) and ProteinMPNN(43)) and outperformed the ESM2 model, despite the fact that TXpredict utilizes ESM2 embeddings as input. TXpredict also showed weaker correlations with predictions from MSA transformer(41) and TranceptEVE(44). A detailed residue-level analysis for THO1 revealed that TXpredict’s expression prediction successfully captures the negative functional impact of an arginine substitution at residue 19 within the SAP domain (Fig. 3e). Interestingly, for the yeast protein GAL4, expression predictions derived from partial mutant sequences correlated more strongly with protein mutation effects than predictions from the full-length sequences (Fig. 3f).

**Figure 3.**
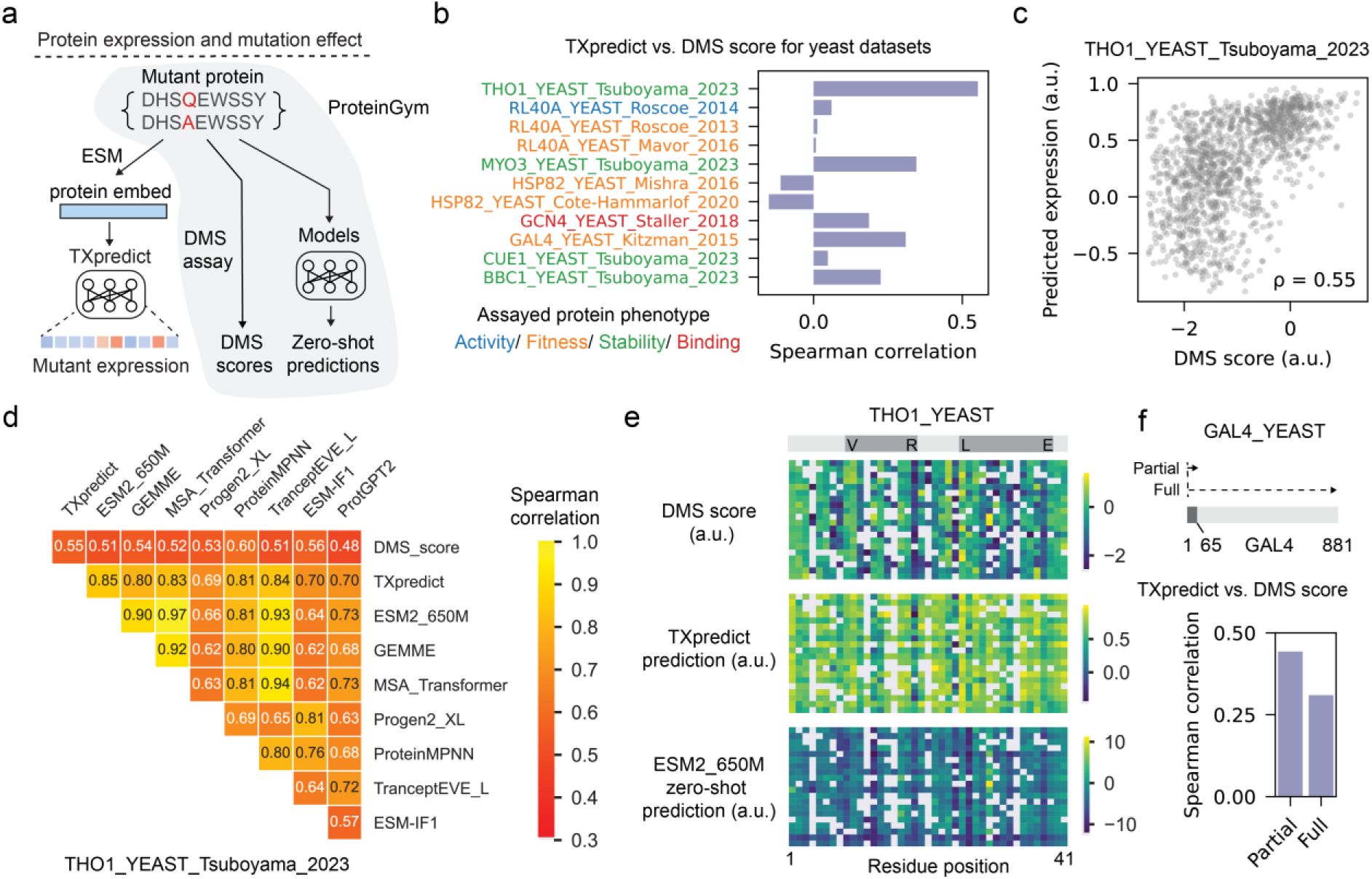
Relationship between protein expression and mutation effects. **a)** Schematic showing relationships among TXpredict predictions, experimentally measured deep mutational scanning (DMS) scores, and zero-shot predictions from ProteinGym30. **b)** Spearman correlation between TXpredict predictions and experimental DMS scores in yeast datasets. **c)** Scattering plot showing predicted expression versus DMS scores for the THO1 protein mutants (n = 1, 279). Spearman correlation coefficient is reported in the bottom-right corner. **d)** Cross-correlations of TXpredict, zero-shot model predictions, and DMS scores for THO1 mutants. **e)** Detailed comparison of DMS scores, TXpredict predictions, and ESM2 predictions across residue positions for single amino acid mutations in THO1. The alpha-helical domains are indicated by dark gray boxes. **f)** Correlation of TXpredict predictions for either a partial region or the full-length GAL4 protein with experimentally measured mutation effects. The mutated part of GAL4 is colored in dark gray.

### Application of TXpredict to new genomes

We applied the trained TXpredict model to predict transcriptomes for new genomes including 2, 169 bacterial, 423 archaeal and 93 fungal species (Fig. 4a and b). Collectively, these new genomes span 1, 744 genera, of which 82% have no prior RNA-seq data according to publicly available data in the NCBI GEO database (Fig. 4 c and d). To explore the relationship between predicted gene expression and protein function, we used DeepFRI(32)(39) to predict Gene Ontology (GO) terms for the new genes. Among the GO terms with the highest predicted expression levels were structural constituents of ribosome (GO:0003735) (Fig. 4e, Supplementary Fig. 19). In contrast, GO terms with the lowest predicted expression included proteins with DNA-binding activity (GO:0003677), consistent with their function as transcriptional regulators. Collectively, our model predicted the expression of up to 9.8M genes across these genomes, which were compiled into a database named TXpredictDB. The predicted gene expressions in different domains followed similar distributions (Fig. 4f). When visualizing protein embeddings of all proteins, we observed a clear separation of bacterial, archaeal and fungal genes (Fig. 4g, Supplementary Fig. 20). Small clusters of ribosomal genes located at the periphery of the plot and showed high predicted expression, while transmembrane transporter proteins formed distinct low-expression clusters.

**Figure 4.**
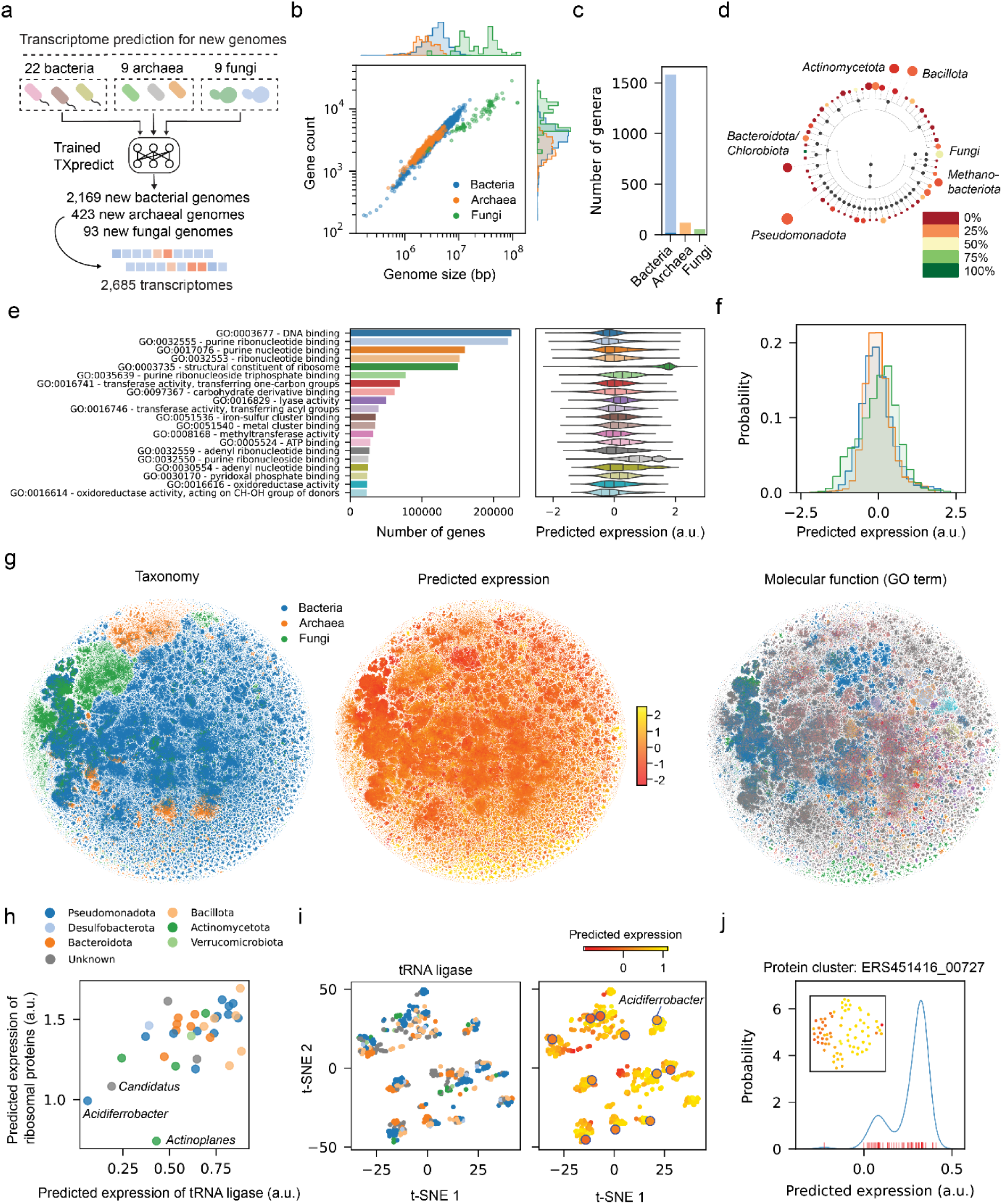
Application of TXpredict to new genomes. **a)** Transcriptome prediction for new genomes. **b)** Genome sizes and gene counts for the new genomes (n = 2, 685). **c)** Number of genera in the new genomes (light colors) compared to training genomes (dark colors). **d)** Polygenetic tree of newly added genomes. Each dot represents a phylum. The dot size is proportional to the number of genera it contains and color indicates the proportion of genera with existing RNA-seq data based on the NCBI GEO database. **e)** Top 20 GO terms of the genes in the new genomes calculated by the DeepFRI model39 (left) and their predicted gene expression levels (right). **f)** Distributions of predicted gene expression levels across new bacterial (blue), archaeal (orange) and fungal (green) genomes. **g)** t-SNE plots of protein embeddings generated by the ESM2 model24 (n = 9.8M). Each dot represents a gene in the new genomes, colored by taxa (left), predicted gene expression (middle), and GO terms (right). GO terms are colored as in panel (**e**). **h)** Predicted expression of tRNA ligases and ribosomal proteins across understudied genera (n = 40), colored by phylum. **i)** t-SNE plots of tRNA ligase embeddings across understudied genera, colored by phylum (left) and predicted expression level (right). **j)** Histogram of predicted expression levels for proteins in a high-similarity cluster. Inset shows the t-SNE plot of embeddings for these proteins, colored by predicted expression.

To explore the transcriptional landscape of understudied species, we focused on genera without prior RNA-seq data and grouped genes based on their annotated Gene Ontology (GO) terms (Supplementary Fig. 21). There was a positive correlation between the predicted expression levels of tRNA ligases and ribosomal proteins across genera (Fig. 4h, Spearman correlation 0.52). *Actinoplanes* had the lowest predicted expression for ribosomal proteins, in accord with their known slow growth rate(48). The endosymbiont Candidatus *Blochmannia*, found in carpenter ants, showed low expression of both tRNA ligases and ribosomal proteins. This pattern likely reflects its reduced need for protein turnover and genome stasis(49), in contrast to free-living bacteria. We further visualized the embeddings of tRNA ligases to examine their sequence variations (Fig. 4i). tRNA ligases from Ca. *Blochmannia* formed distinct low-expression clusters, whereas those from *Acidiferrobacter* were distributed across multiple clusters despite similarly low expression levels, suggesting divergent evolutionary constraints associated with their respective ecological niches..

We also clustered protein sequences using MMseqs2(33) and predicted gene expression within each high-similarity cluster using the TXpredict model. Interestingly, several clusters displayed non-unimodal expression distributions, with conserved amino acid differences separating low- and high-expression variants (Fig. 4j). For example, in cluster of the PTS IIA component homologs, low-expression variants shared a V116K substitution (Supplementary Fig. 22), which could reduce translation efficiency and mRNA stability due to the presence of constitutive lysine residues(50, 51).

We further analyzed the predicted gene expression profiles of biosynthetic gene clusters (BGCs) across bacterial species in TXpredictDB, which were identified using antiSMASH(52)(45) (n = 11, 400). Although BGCs are typically lowly expressed, distinct patterns were observed across different cluster types. For example, the proteusin BGC showed high expression of core biosynthetic genes, whereas the ectoine BGC exhibited elevated expression of regulatory genes (Supplementary Fig. 23). These reference expression profiles in non-model organisms may offer valuable insights into the functional organization of BGCs.

## DISCUSSION

We present TXpredict, a transcriptome prediction model that can generalize to a wide range of microbes. For a microbial genome with 4.6k genes, transcriptome prediction can be completed in approximately 22 minutes using a Google Colab notebook (Material and Methods), demonstrating the model’s efficiency and accessibility. Our approach is not without limitations: while it effectively leverages protein embeddings for expression prediction, it does not account for the effects of cis-regulatory elements. Moreover, our model’s performance is constrained by the limited training profiles, especially for archaeal species. Lastly, we provide a single reference transcriptome per genome in TXpredictDB, which does not capture gene expression dynamics. Our condition-dependent model captures context-dependent expression patterns for unseen genes, but it does not extrapolate to entirely novel experimental conditions not represented in the training set.

Our training data for TXpredict relies on next-generation sequencing (NGS) data from GEO datasets, as they currently provide the most comprehensive coverage of bacterial species and experimental conditions available. Nanopore-based direct RNA sequencing offers several advantages, including full-length transcript detection and the avoidance of amplification and fragmentation artifacts inherent to short-read methodologies(28–30, 53–57). While our model captures platform-independent biological signals, we anticipate that incorporating additional nanopore datasets as they become available will further improve the model accuracy.

Despite these limitations, we propose several potential directions to further improve our model’s performance. First, employing more powerful pre-trained language models could produce richer embeddings that better capture evolutionary constraints relevant to expression prediction. Moreover, integrating multi-modal sequence inputs (protein, DNA, and codon-aware representations) may provide complementary signals at different levels. Third, for data-sparse species, structure- and homology-aware augmentation or transfer learning method could potentially reduce model bias and improve its generalization. Finally, including more training species and broader condition coverage would further enhance performance.

TXpredict has the potential to complement experimental approaches and advance the study of the vast number of sequenced yet poorly characterized microbes. For example, identifying conserved expressed genes in pathogens may enable the discovery of novel drug targets(58). Furthermore, predicting gene expression across genomes can provide insight into how genome plasticity contributes to bacterial pathogenicity(59). Our model may also suggest biologically plausible mRNA expression levels as starting points for heterologous gene expression in synthetic biology applications. These efforts could further benefit from integrating single-cell RNA-seq data(5, 60, 61) and computational tools for mRNA synthesis and stability prediction(62–64), paving the way for a deeper understanding of microbial transcriptional landscapes.

## DATA AVAILABILITY

Our trained model and inference codes are available from GitHub: https://github.com/lingxusb/TXpredict

