## Supplementary Information for "Predicting microbial transcriptome using annotated genome sequence"

**SUPPLEMENTARY DATA**

**Supplementary Table S1.** Datasets used in the transcriptome prediction model.

| **Bacteria** | | | | |
|---|---|---|---|---|
| **Species** | **GSE number** | **Reference genome** | **Gene#** | **Condition #** |
| *Bacillus subtilis* | GSE224332(65) | NZ_CP020102.1 | 4220 | 317 |
| *Bacteroides thetaiotaomicron* | GSE251676(66) | GCF_000011065.1 | 4618 | 64 |
| *Bordetella hinzii* | GSE181542 | CP052845.1 | 4531 | 228 |
| *Caulobacter vibrioides* | GSE246782(67) | NC_011916.1 | 3882 | 34 |
| *Cereibacter sphaeroides* | GSE186600(68) | GCA_000012905.2 | 4232 | 17 |
| *Chlamydia trachomatis* | GSE262652 | GCA_000068585.1 | 857 | 30 |
| *Clostridioides difficile* | GSE249810(69) | GCA_000009205.2 | 3560 | 24 |
| *Clostridium acetobutylicum* | GSE107804(70) | GCA_000008765.1 | 3755 | 48 |
| *Clostridium beijerinckii* | GSE150480(71) | GCF_000833105.2 | 5316 | 16 |
| *Clostridium butyricum* | GSE150480(71) | GCF_001456065.2 | 3848 | 16 |
| *Clostridium tetani* | GSE182408(72) | NC_004557.1 | 2562 | 39 |
| *Escherichia coli* | GSE152664(73) | NC_010473.1 | 3260 | 9 |
| *Erythrobacter litoralis* | GSE126532 | GCA_001719165.1 | 3003 | 49 |
| *Methylotuvimicrobium buryatense* | GSE162089(74) | NZ_CP035467.1 | 3927 | 56 |
| *Mycobacterium tuberculosis* | GSE110508(75) | GCA_000195955.2 | 3957 | 87 |
| *Mycolicibacterium smegmatis* | GSE69983(76) | GCA_000015005.1 | 6700 | 30 |

| | | | | |
|---|---|---|---|---|
| *Pseudomonas aeruginosa* | GSE251671 | GCA_000014625.1 | 5659 | 858 |
| *Pseudomonas syringae* | GSE103441(77) | GCA_000007805.1 | 5586 | 114 |
| *Streptococcus dysgalactiae* | GSE272047(78) | GCA_029234115.1 | 1939 | 114 |
| *Streptococcus pyogenes* | GSE113055(79) | CP000056.2 | 1608 | 442 |
| *Xanthomonas citri* | GSE252662(80) | AE008923.1 | 4294 | 38 |
| *Zymomonas mobilis* | GSE139939(81) | CP023715.1 | 1759 | 19 |
| **Summary** | | | 83,073 | 2,649 |

Supplementary Table S1 - continue. Datasets used in the transcriptome prediction model.

| **Archaea** | | | | |
|---|---|---|---|---|
| **Species** | **GSE number** | **Reference genome** | **Gene#** | **Condition#** |
| *Haloarcula hispanica* | GSE227031 | GCF_000223905.1 | 3776 | 32 |
| *Halobacterium salinarum* | GSE182493(82) | GCF_000006805.1 | 2325 | 24 |
| *Haloferax volcanii* | GSE248564(83) | GCF_000025685.1 | 3873 | 39 |
| *Methanocaldococcus jannaschii* | GSE112986(84) | GCF_000091665.1 | 1724 | 13 |
| *Methanococcus maripaludis* | GSE261517(85) | GCF_000011585.1 | 1730 | 12 |
| *Methanosarcina barkeri* | GSE168895(86) | GCF_000970025.1 | 3338 | 12 |
| *Sulfolobus islandicus* | GSE220819(87) | GCF_000189555.1 | 2682 | 18 |
| *Thermococcus kodakarensis* | GSE186817 | NC_006624.1 | 2214 | 4 |
| *Thermococcus paralvinellae* | GSE101078(88) | GCF_000517445.1 | 1995 | 15 |
| **Summary** | | | 25,448 | 169 |

Supplementary Table S1 - continue. Datasets used in the transcriptome prediction model.

| **Fungi** | | | | |
|---|---|---|---|---|
| **Species** | **GSE number** | **Reference genome** | **Gene#** | **Conditio#** |
| *Clavispora lusitaniae* | GSE162151(89) | ATCC42720_w_ CBS_6936_MT | 5851 | 54 |
| *Cryptococcus neoformans* | GSE226255(90) | CP022321.1 | 6718 | 720 |
| *Saccharomyces cerevisiae* | GSE234431 | GCA_000146045.2 | 5498 | 897 |
| *Coccidioides posadasii* | GSE279406 | NC_089407.1 | 8103 | 144 |
| *Blumeria graminis* | GSE69215(91) | GCA_000151065.3 | 6329 | 48 |
| *Lipomyces starkeyi* | GSE276000 | GCA_001661325.1 | 7928 | 36 |
| *Neurospora crassa* | GSE181566 | GCF_000182925.2 | 9532 | 80 |
| *Candida albicans* | GSE159545(92) | GCF_000182965.3 | 5918 | 75 |
| *Orbilia oligospora* | GSE245014 | GCF_000225545.1 | 11195 | 24 |
| **Summary** | | | 67,072 | 2,078 |

**Supplementary Table S2.** Datasets used in the conditional gene expression model (https://imodulondb.org/)

| Species | Reference | Reference genome | Gene# | Condition # |
|---|---|---|---|---|
| *Acinetobacter baumannii* | Menon, et al.(93) | CP008706.1 | 1788 | 139 |
| *Bacillus subtilis* | Sastry, et al. | NC_000964.3 | 1969 | 265 |
| *Escherichia coli* | Lamoureux, et al.(94) | U00096.3 | 2490 | 1035 |
| *Limosilactobacillus reuteri* | Josephs-Spaulding, et al.(95) | GCA_020785475.1 | 1159 | 117 |
| *Mycobacterium tuberculosis* | Yoo, et al.(96) | AL123456.3 | 3214 | 647 |
| *Pseudomonas aeruginosa* | Rajput, et al.(97) | NC_002516.2 | 3981 | 411 |
| *Pseudomonas putida* | Lim, et al.(1) | AE015451.2 | 4379 | 321 |
| *Pseudomonas syringae* | Bajpe, et al.(98) | AE016853.1 | 4296 | 202 |
| *Staphylococcus aureus* | Poudel, et al.(99) | NC_010079.1 | 1736 | 385 |
| *Synechococcus elongatus* | Yuan, et al.(100) | NC_007604.1 | 1522 | 300 |
| *Salmonella enterica* | Yuan, et al.(100) | AE006468.2 | 1808 | 533 |
| *Streptococcus pyogenes* | Hirose, et al.(101) | CP008776.1 | 1173 | 116 |
| *Vibrio natriegens* | Shin, et al.(4) | GCA_001456255.1 | 1981 | 148 |
| **Summary** | | | 31,496 | 4,619 |

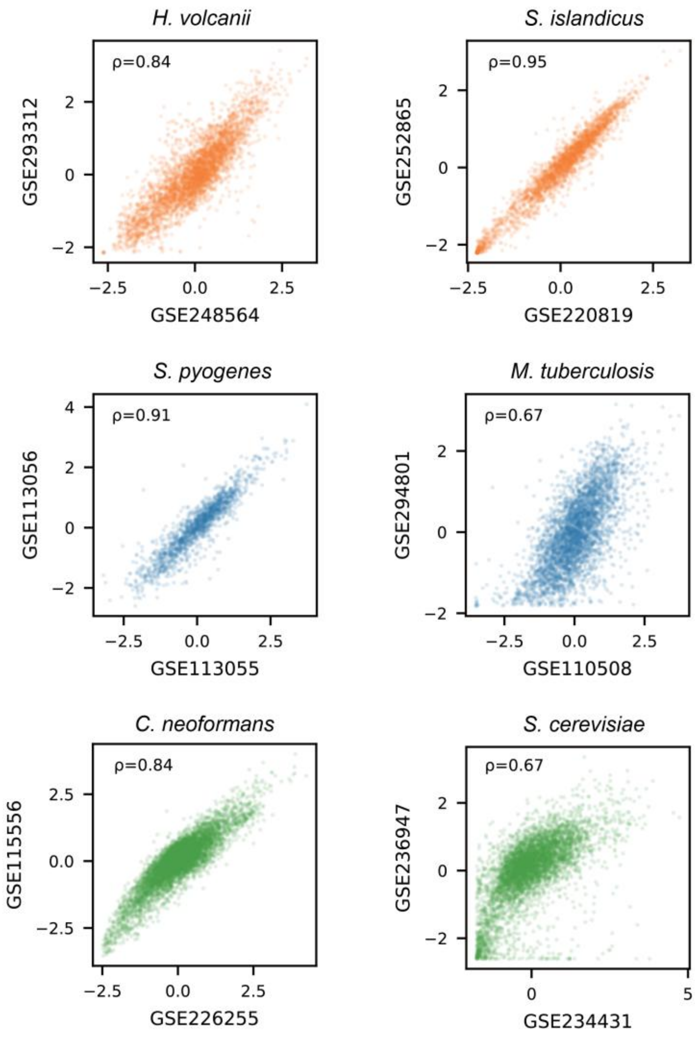


**Supplementary Figure 1.** Comparison of mean gene expression across GEO datasets for the same species. The Spearman correlation coefficients of two dataset's mean gene expression are reported in the top-left corner of each plot. The GEO dataset IDs are shown as axis labels.

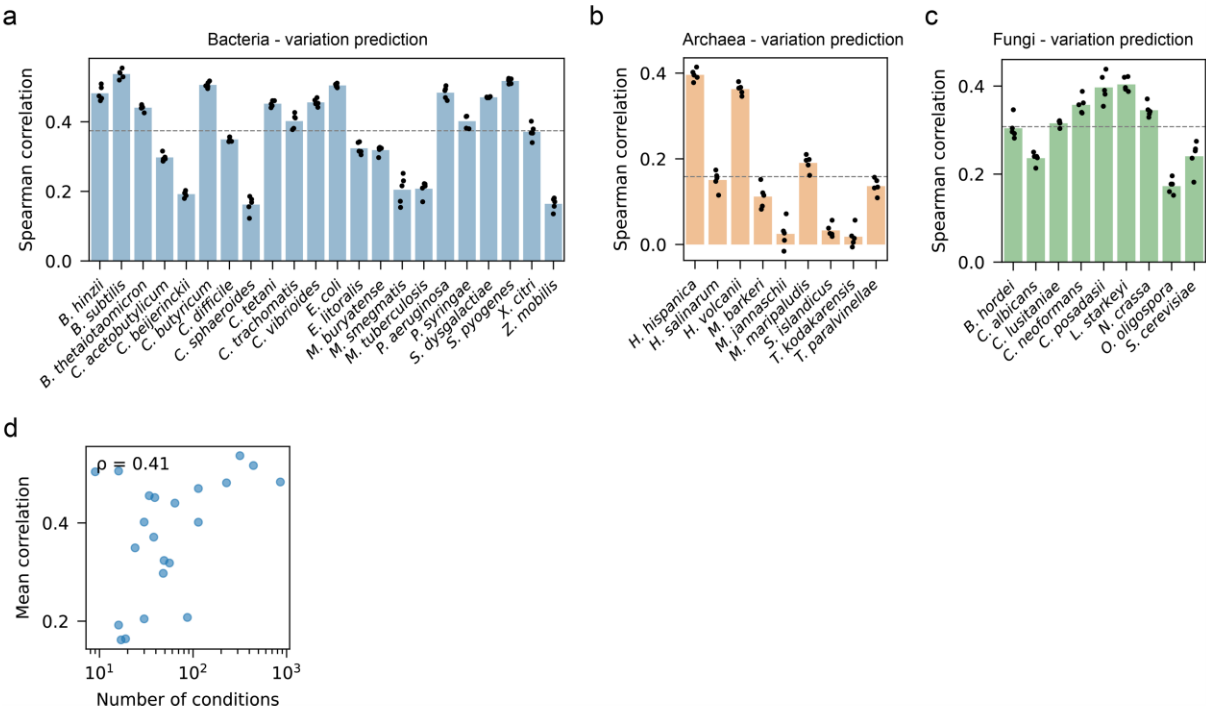


**Supplementary Figure 2.** Prediction of gene expression variation. Spearman correlations between predicted and measured gene expression variations in leave-one-genome-out experiments for bacterial **(a)**, archaeal **(b)**, and fungal **(c)** species. Results from five rounds of model training and evaluation are shown (n = 5). **d)** Number of conditions and corresponding Spearman correlations for expression variation prediction in bacterial species (n = 22).

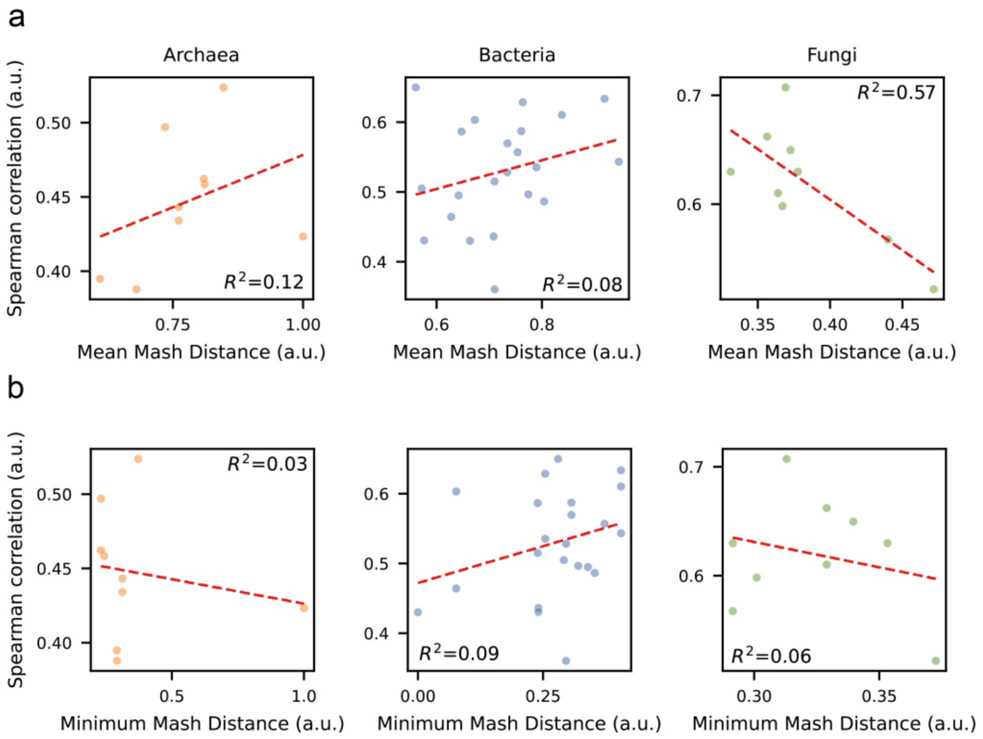


**Supplementary Figure 3.** Evolutionary distance and Spearman correlation for each species under leave-one-genome-out evaluation. **a)** Scatter plot of the mean Mash distance between the test genome and training genomes versus the correlation between true and predicted gene expression for the test genome. The dashed red line shows the linear fit with $R^2$ . **b)** Scatter plot of the minimum Mash distance between the test genome and training genomes versus the correlation between true and predicted gene expression for the test genome. The dashed red line shows the linear fit with $R^2$.

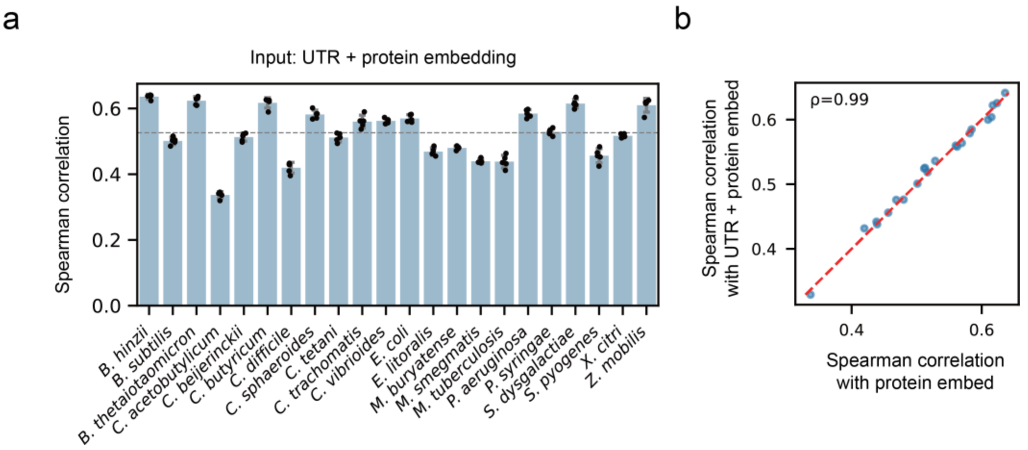


**Supplementary Figure 4.** Transcriptome model performance using UTR + protein sequences on bacterial datasets. **a)** Spearman correlations between predicted and measured gene expression in leave-one-genome-out evaluations with UTR + protein embeddings. Dashed line indicates the mean correlations for all bacterial datasets (0.53). **b)** Scatter plot comparing correlations from different inputs (UTR + protein vs protein-only). The red dashed line represents y=x.

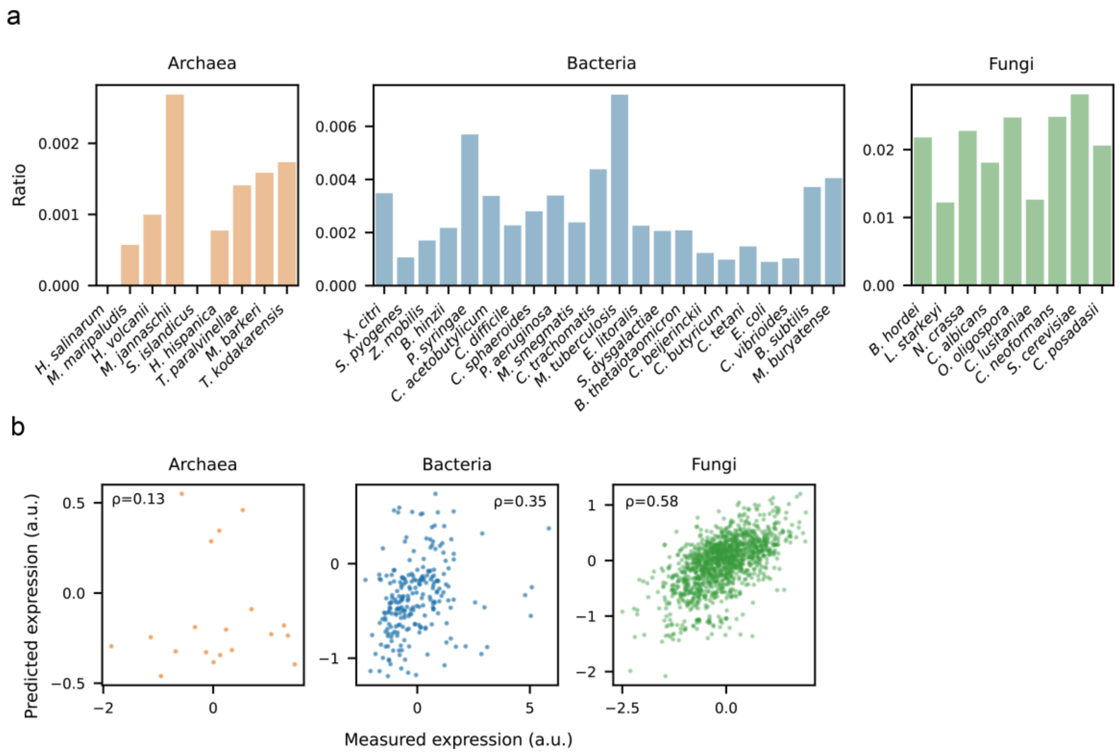


**Supplementary Figure 5.** Analysis of proteins longer than 1,500 amino acids. **a)** The ratio of proteins longer than 1500 amino acids in each species. **b)** Scatter plots comparing TXpredict and measured expression for proteins longer than 1500 AA. The Spearman correlation coefficients are reported in the corner of each plot.

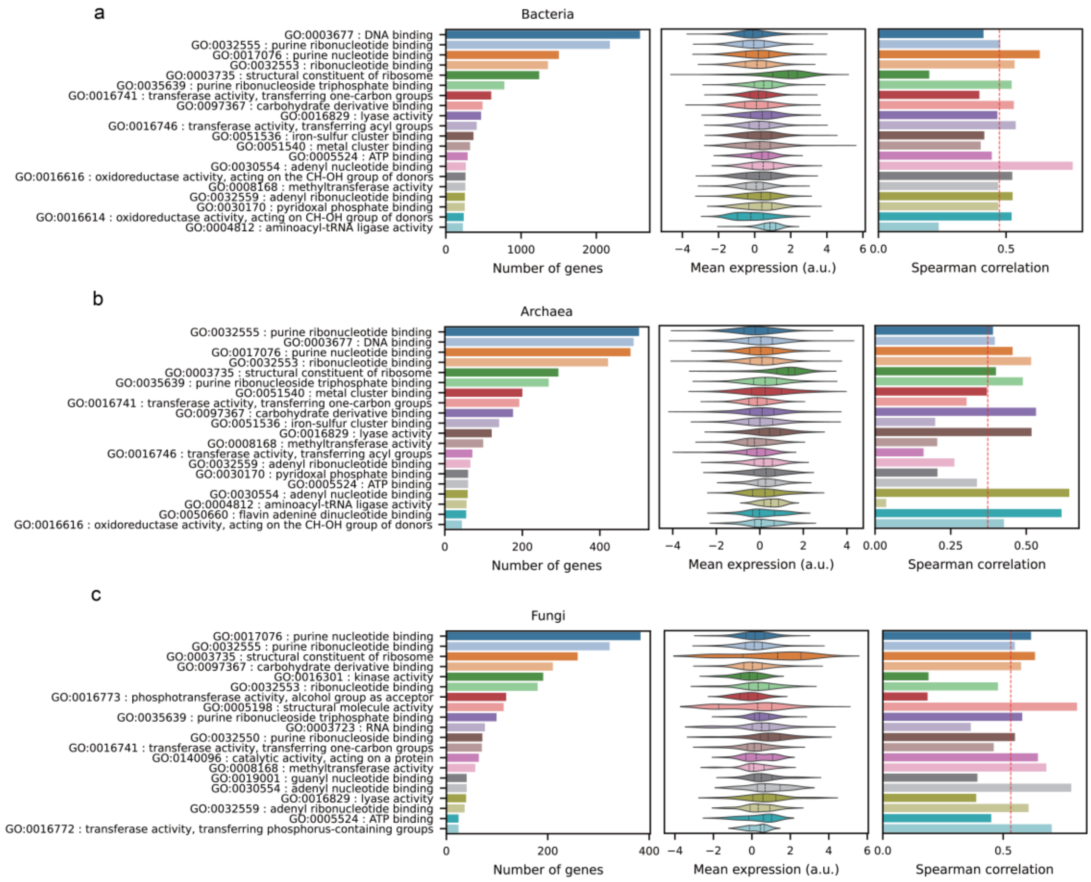


**Supplementary Figure 6.** Top GO terms for the test genes in the leave-one-genome-out cross validation. For each GO term, we report the number of genes, mean expression, and Spearman correlations between measured and predicted gene expression. **a)** Result of bacteria. The red dashed line represents the top GO terms' mean spearman correlation (0.47). **b)** Result of archaea. The red dashed line represents the top GO terms' mean spearman correlation (0.37). **c)** Result of fungi. The red dashed line represents the top GO terms' mean spearman correlation (0.53).

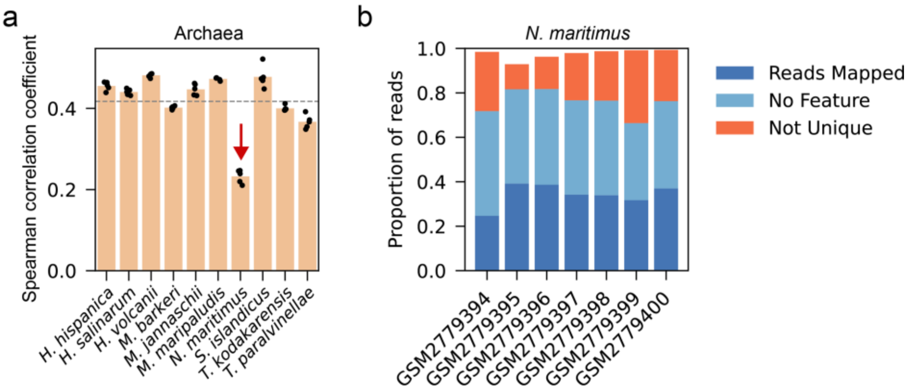


**Supplementary Figure 7.** Detection of issues related to RNA-seq dataset. **a)** Spearman correlation coefficients between predicted and measured gene expression for all archaeal species in the initial evaluation. Results from five rounds of model training and testing are shown (n = 5). **b)** Proportion of mapped/unmapped reads in the *N. maritimus* dataset (GSE103699).

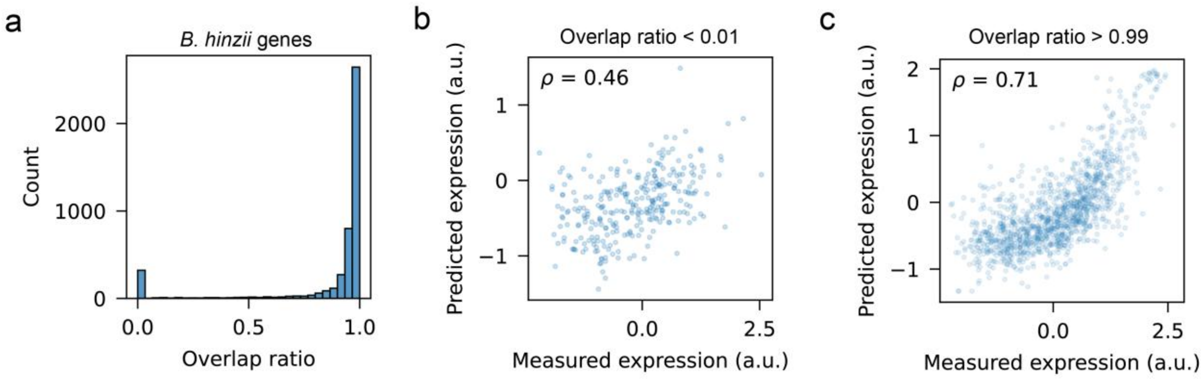


**Supplementary Figure 8.** Expression prediction for rare proteins. **a)** Histogram showing the distribution of overlap ratios between *B. hinzii* proteins and proteins in the training dataset. Overlap ratios were calculated as the aligned sequence length divided by the query length. **b)** Predicted versus measured gene expression for proteins with low homology to the training set (overlap ratio < 0.01). **c)** Predicted and measured gene expressions for proteins with close homologs in the training dataset (overlap > 0.99).

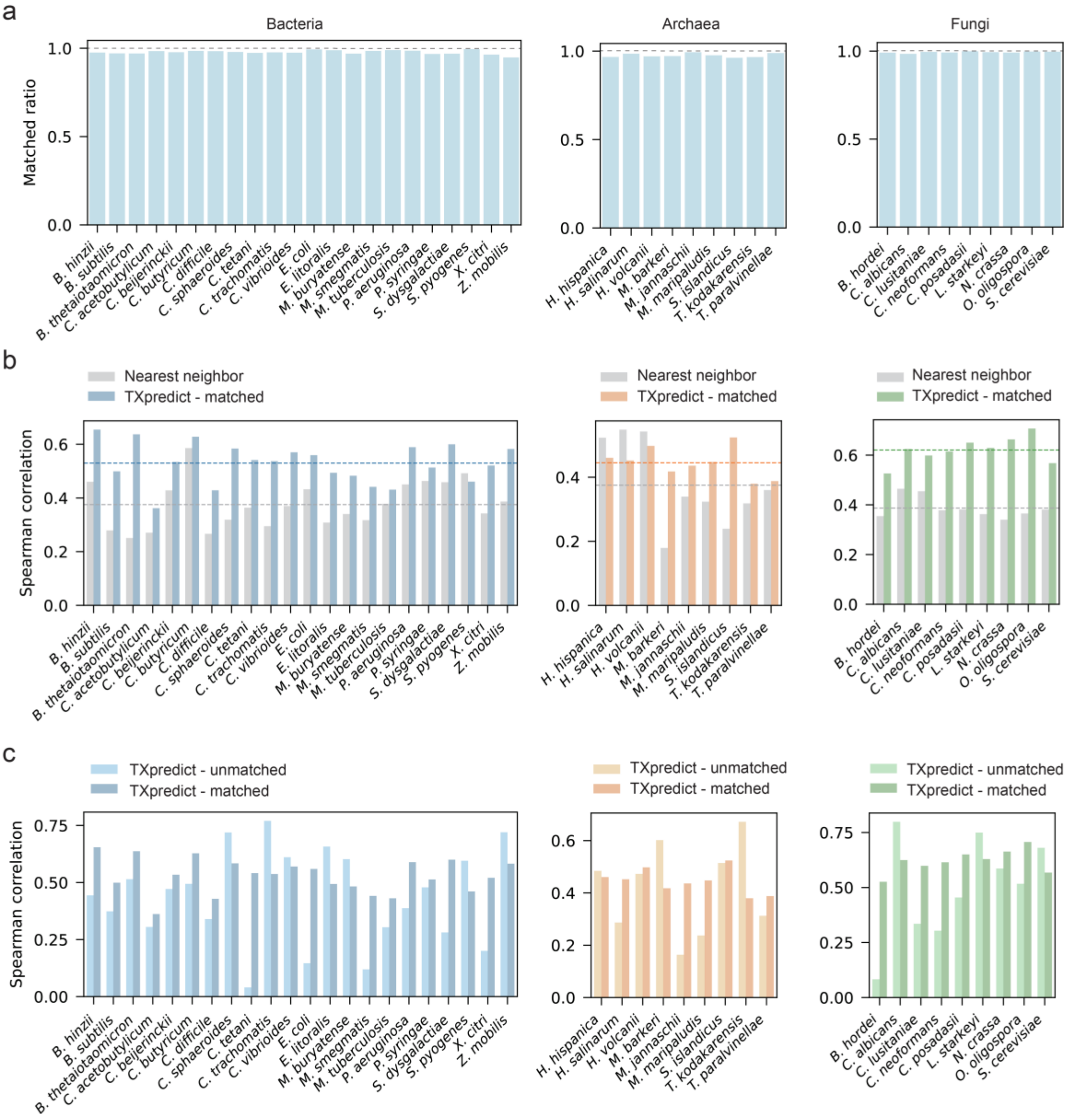


**Supplementary Figure 9.** Benchmarking the performance of the TXpredict model against the one-nearest-neighbor method. **a)** Proportion of proteins in the test genome that have a BLASTP match in the training data. **b)** Spearman correlations between true and predicted gene expression for the one-nearest-neighbor baseline and TXpredict under leave-one-genome-out evaluation. TXpredict's mean correlations for bacteria, archaea and fungi is respectively 0.53, 0.44 and 0.62, compared to the BLASTP-based baseline values of 0.38, 0.37 and 0.39. Dashed lines denote the mean correlation of each method within each domain. **c)** Spearman correlations between predictions and mean expression for TXpredict, comparing BLASTP-matched and BLASTP-unmatched proteins within each species. The mean correlations for unmatched proteins of bacteria, archaea and fungi are 0.44, 0.42 and 0.50 respectively.

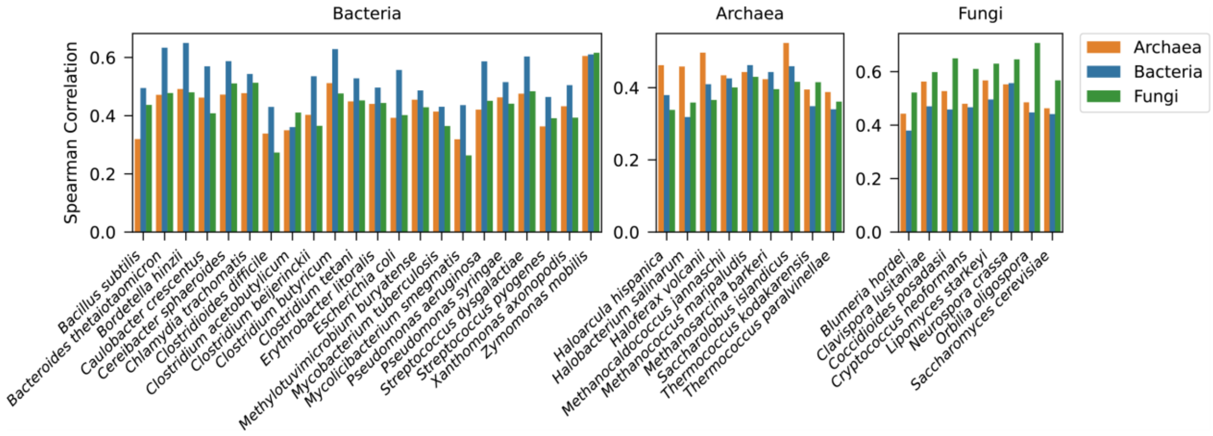


**Supplementary Figure 10.** Cross-domain prediction performance of TXpredict. TXpredict was trained on all species from a single domain and evaluated on species from the two other domains. Results from three domain-specific models are shown: bacteria (blue), archaea (orange), and fungi (green).

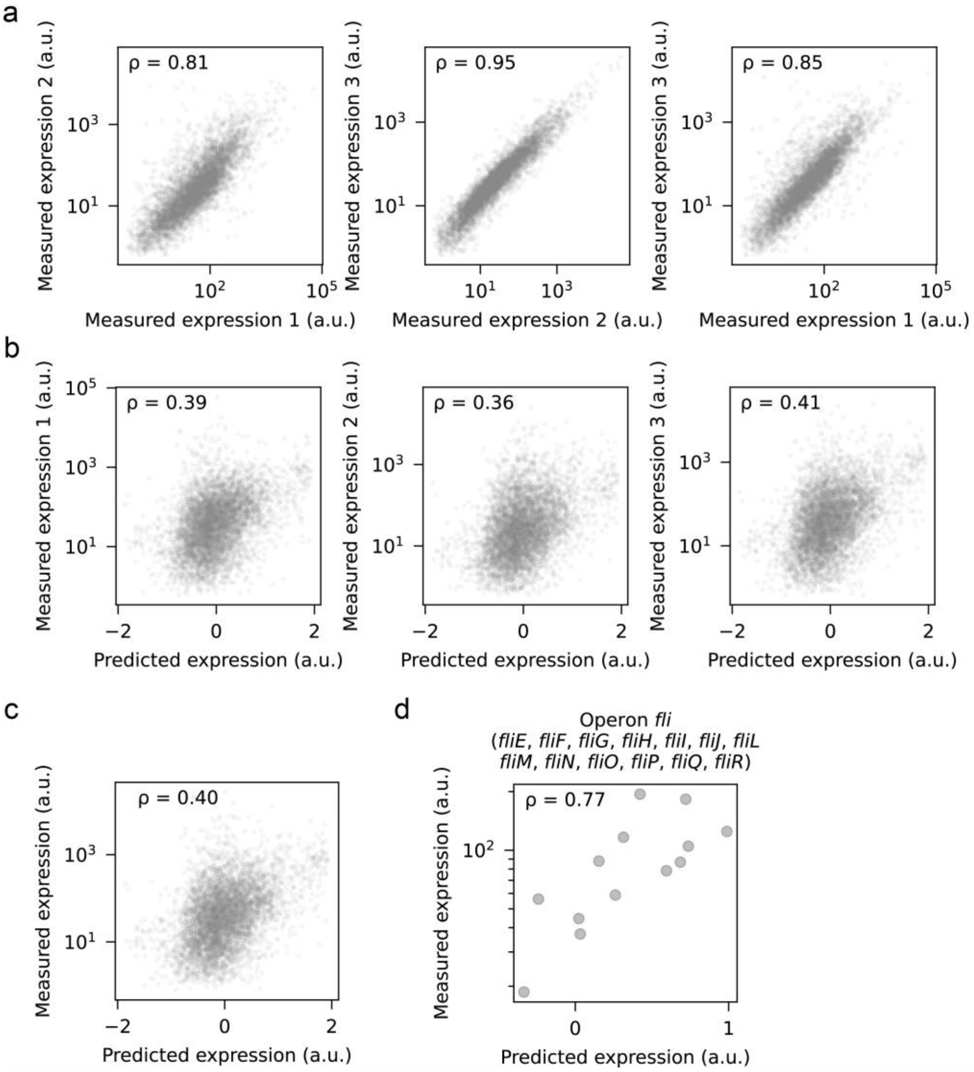


**Supplementary Figure 11.** Performance of TXpredict on *Pseudomonas sp.* WBC-3. **a)** Correlation of measured gene expression values of *Pseudomonas sp. WBC-3* across three experimental conditions. **b)** Predicted versus measured gene expression under each condition. **c)** Scatter plot comparing predicted and mean measured gene expression across three conditions for all genes. **d)** Predicted and measured gene expression levels for genes in the *fli* operon.

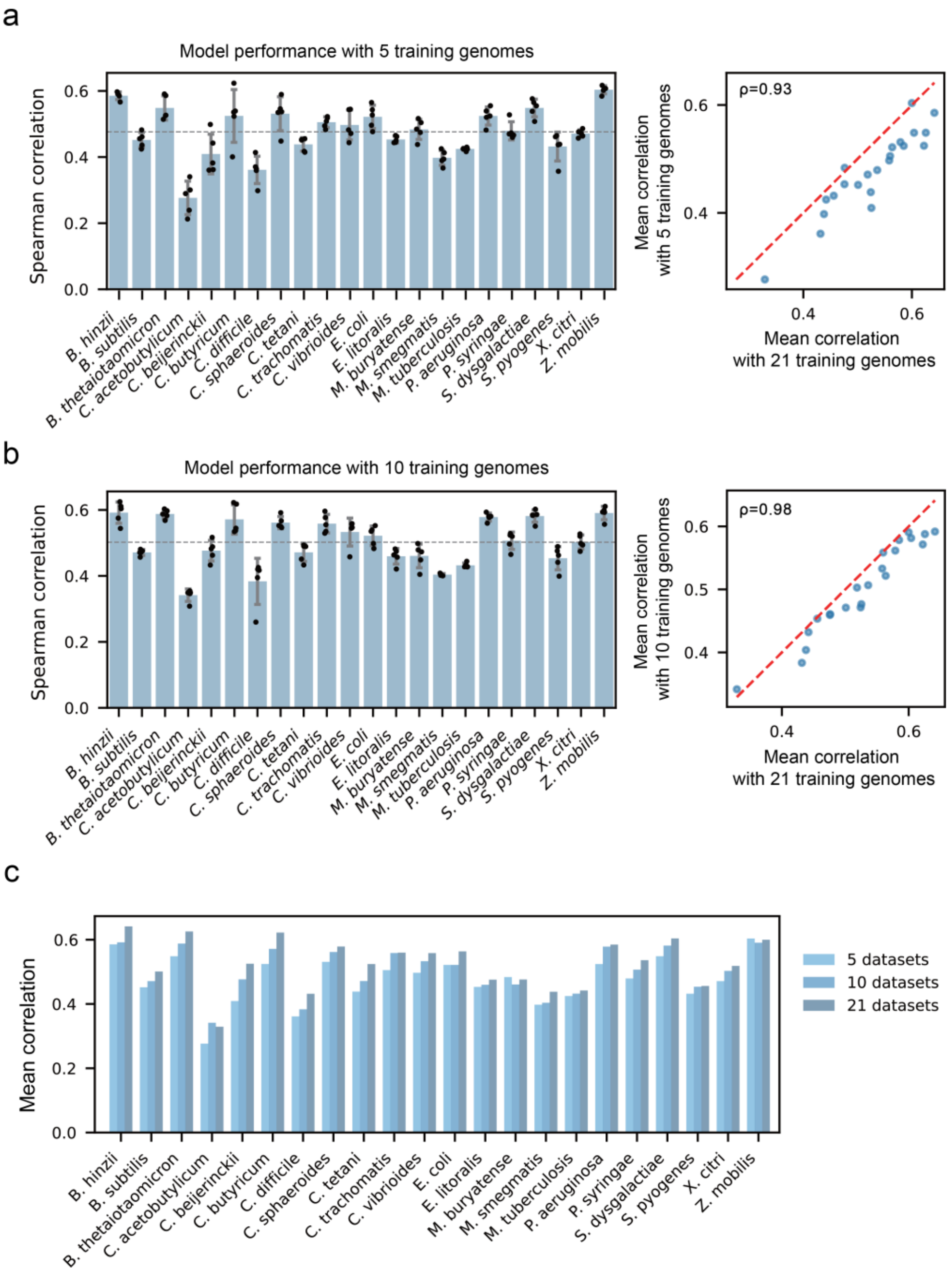


**Supplementary Figure 12.** Scaling analysis of TXpredict performance with varying training data sizes. **a)** Spearman correlations between predicted and measured gene expression in leave-one-genome-out evaluations using 5 randomly selected training genomes. Results from five independent runs are shown (n = 5). Right, scatter plot comparing correlations from 5 training genomes versus all 21 training genomes. The red dashed line represents y=x. **b)** Spearman correlations under the same protocol using 10 randomly selected training genomes (n = 5). Right, scatter plot comparing correlations from 10 training genomes versus all 21 training genomes. The red dashed line represents y=x. **c)** Spearman correlation coefficients between predicted and measured gene expression in leave-one-genome-out evaluations using 5, 10 and all (21) training genomes.

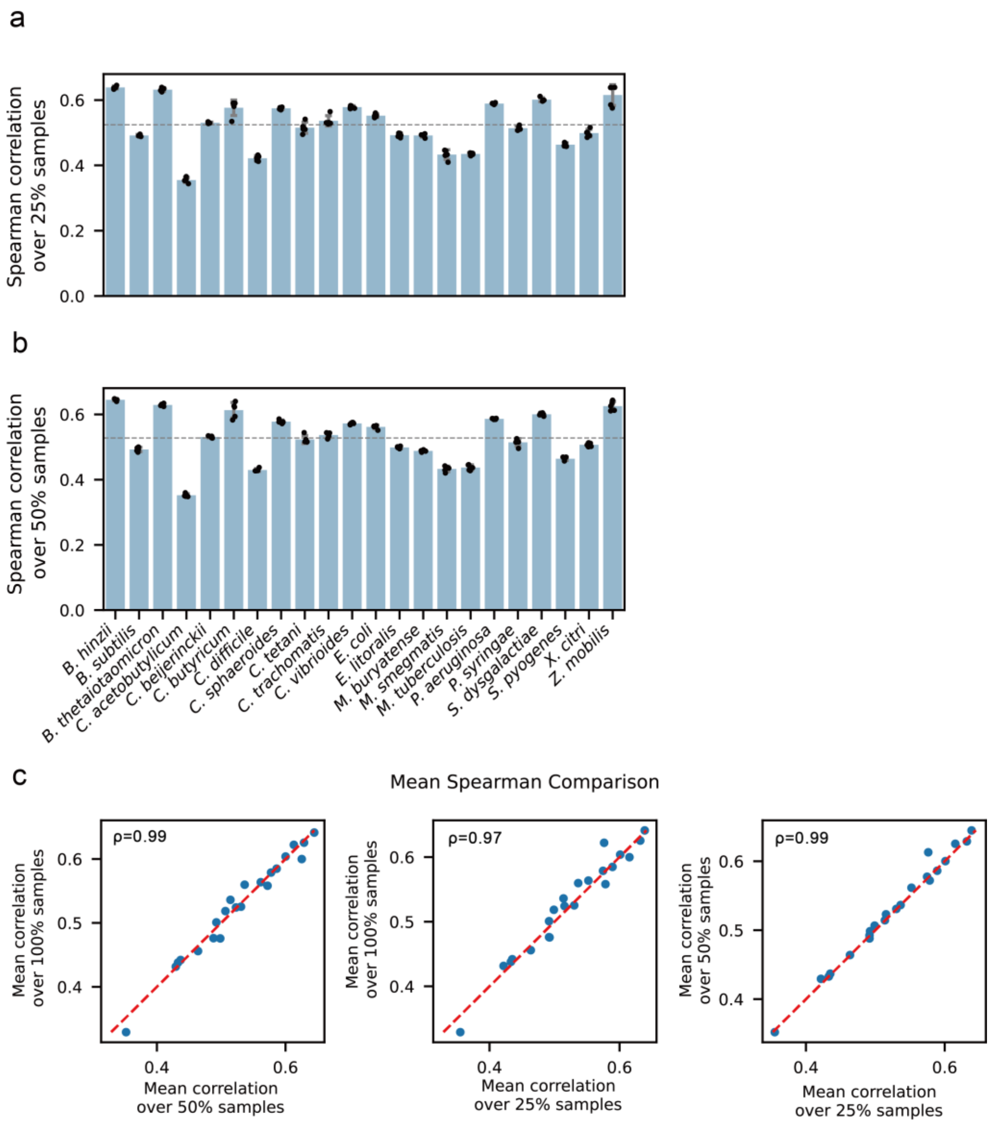


**Supplementary Figure 13.** Subsampling analysis of TXpredict performance. For each bacterial strain, we randomly subsampled conditions to 25% or 50% of all conditions, recomputed mean expression for each gene, and evaluated model performance under leave-one-genome-out. **a)** 25% subsampling. Spearman correlations for each strain across five independent training/testing runs (n = 5). **b)** 50% subsampling. **c)** Scatter plot comparing performance at 25%, 50%, and all conditions. Each point denotes one bacterial strain, and the red dashed line indicates y = x.

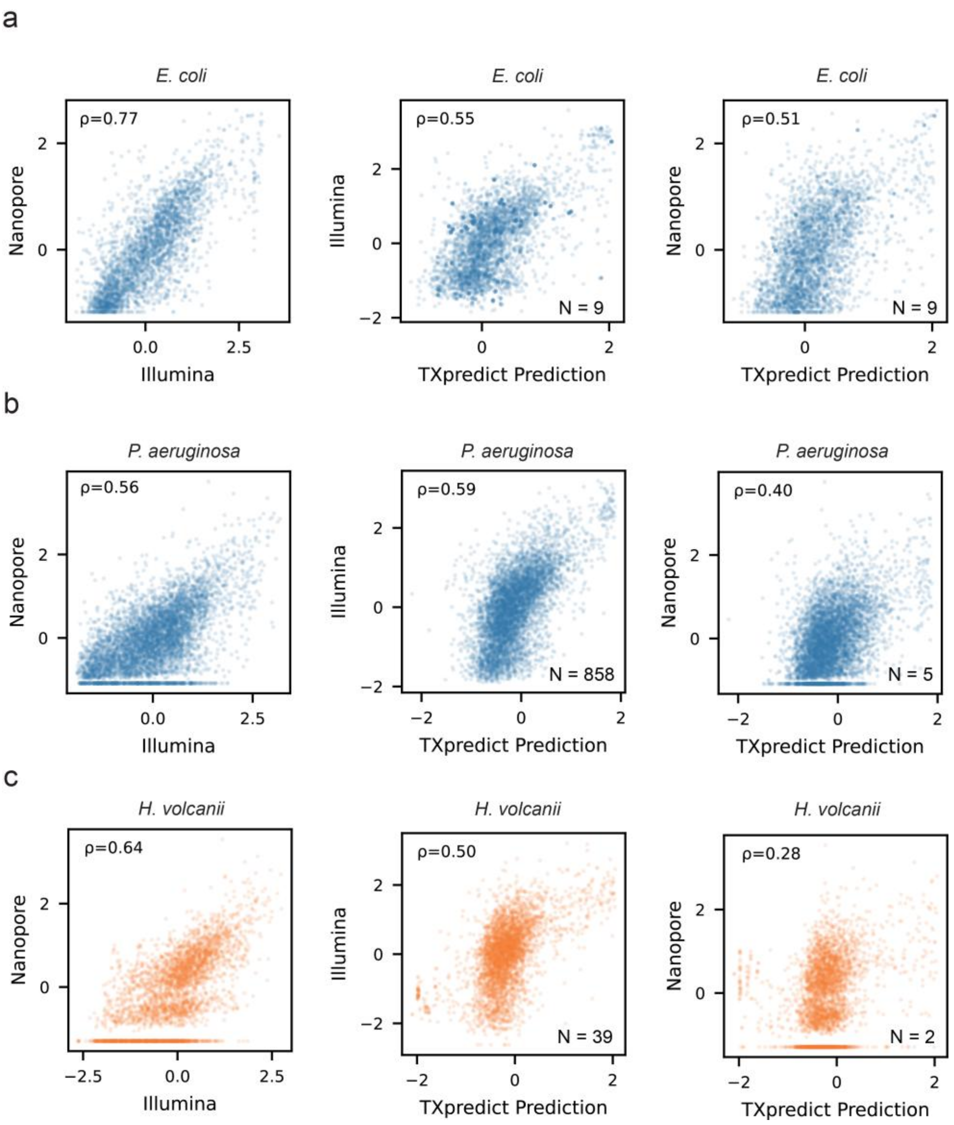


**Supplementary Figure 14.** Comparison of mean gene expression from GEO datasets sequenced with Nanopore or Illumina and TXpredict's predictions. For each species, three scatter plots are shown: Nanopore vs. Illumina; TXpredict vs. Illumina; TXpredict vs. Nanopore. Each dot denotes expression of one gene. Spearman correlations are reported in the top-left of each plot. **a)** *E. coli*. **b)** *P. aeruginosa*. **c)** *H. volcanii*. Sample counts for the Illumina and Nanopore datasets are shown in the lower right corner.

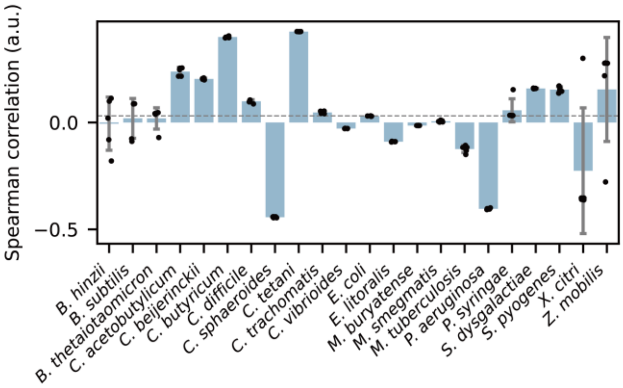


**Supplementary Figure 15.** Spearman correlations between predicted and measured gene expression in leave-one-genome-out evaluations on bacterial species using CodonTransformer embeddings. Results from five independent training/testing runs are shown (n = 5). The dashed line denotes the mean correlation.

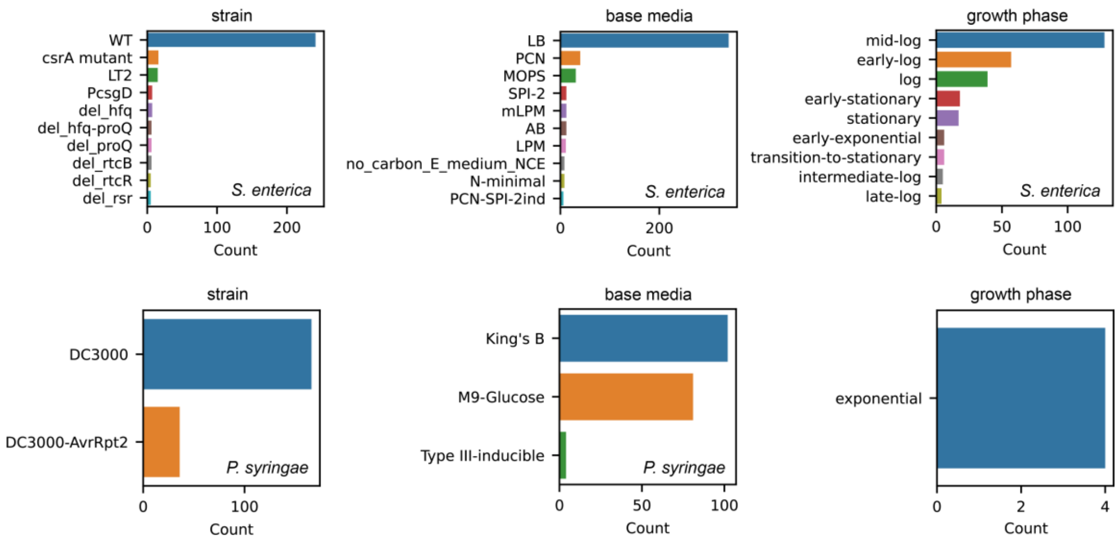


**Supplementary Figure 16.** Diversity of experimental conditions for *S. enterica* and *P. syringae*. The number of conditions were summarized from the sample sheet from iModulonDB.

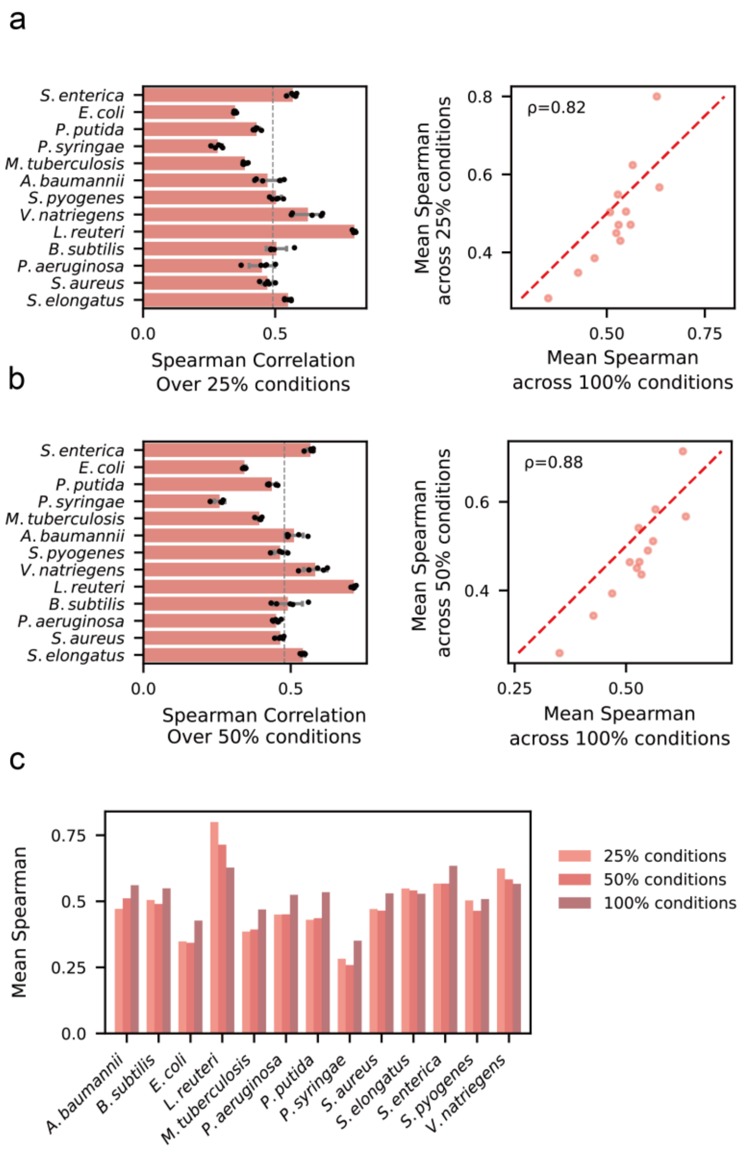


**Supplementary Figure 17.** Impact of experimental condition coverage on the performance of the condition-dependent model. **a)** Spearman correlations between predicted and measured gene expression under 5-fold cross-validation when training on 25% of conditions. Results from five rounds of model training and testing are reported (n = 5). Dashed line indicates the mean correlation (0.49) for all species. Right: scatter plot comparing per-species correlations at 25% vs 100% coverage. The red dashed line represents y=x. **b)** Spearman correlations between predicted and measured gene expression under 5-fold cross-validation when training on 50% of conditions. **c)** Spearman correlation coefficients between predicted and measured gene expression in 5-fold evaluation of condition-dependent models trained on different condition coverages.

a

b

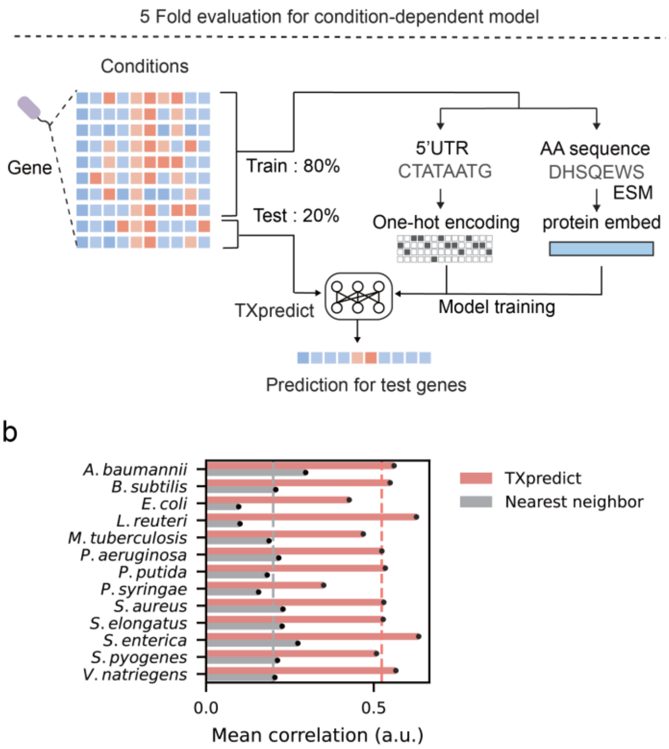


**Supplementary Figure 18.** Benchmarking the condition-dependent model against the one-nearest-neighbor baseline. **a)** Schematic of TXpredict's condition-dependent model under 5-fold cross-validation. **b)** Mean correlation for the BLASTP-based one-nearest-neighbor method and TXpredict across 5 folds. In the nearest neighbor method, each test gene is matched to its closest training gene by BLASTP, and the neighbor's expression is used as the prediction. Dashed lines indicate mean correlation for each method.

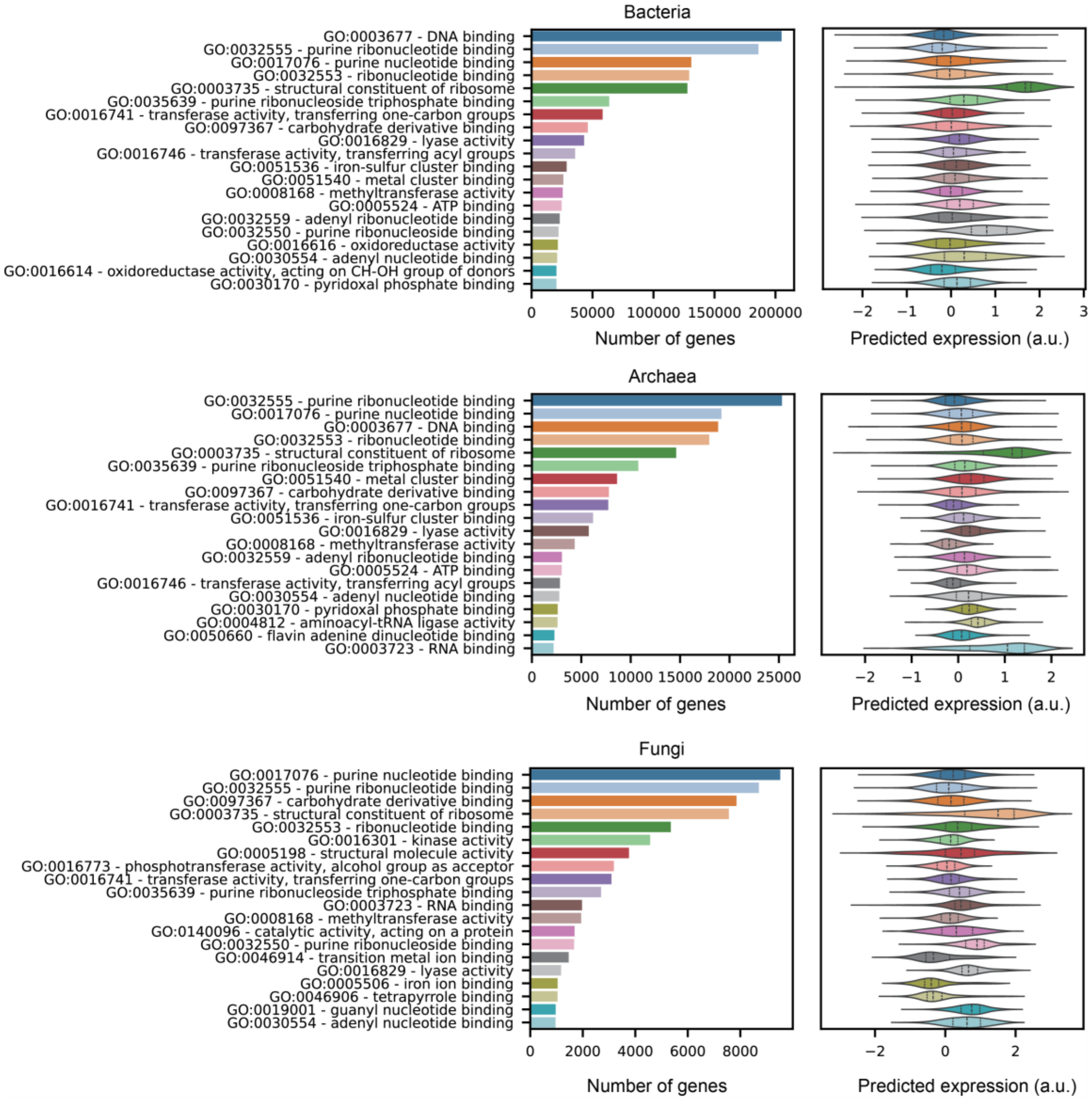


**Supplementary Figure 19.** Top GO terms and the predicted gene expressions in TXpredictDB. Top 20 Gene Ontology (GO) terms for genes from newly added bacterial, archaeal, and fungal genomes, as annotated by the DeepFRI model(39). Predicted gene expression levels for these genes are shown alongside their associated GO terms.

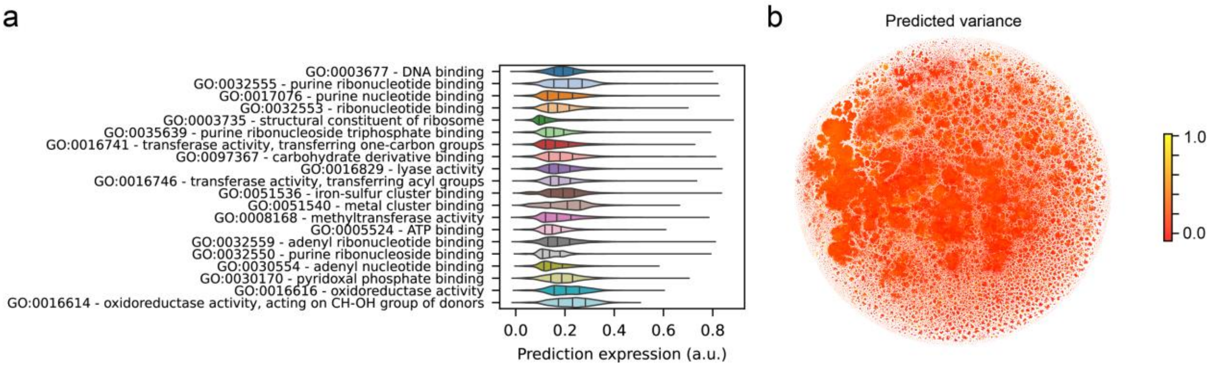


**Supplementary Figure 20.** Prediction of gene expression variation in newly added genomes. **a)** Predicted gene expression variation for genes associated with the top 20 GO terms in the newly added genomes. **b)** Predicted gene expression variation visualized in the t-SNE plot of protein embeddings (n = 9.8M).

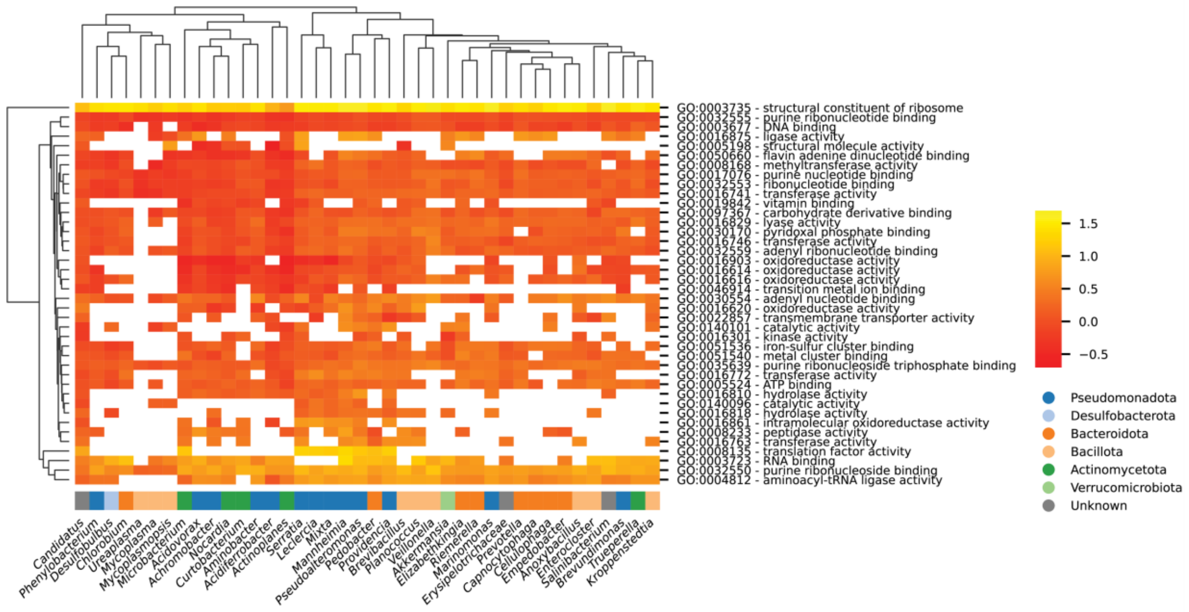


**Supplementary Figure 21.** Prediction of gene expression for understudied genera. Gene expression values for the top 40 GO terms and 40 genera are shown. Only genus–GO term combinations with more than 10 associated genes are included, with mean gene expression levels presented. Genera are colored by phylum.

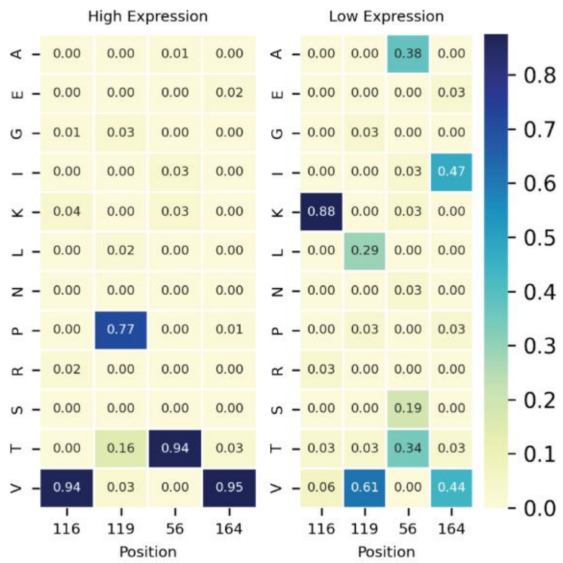


**Supplementary Figure 22.** Amino acid frequency differences between low and high expression genes within a protein cluster. TXpredict was used to predict gene expression for proteins in cluster ERS451416_00727, identified using MMseqs2(42) at a 90% sequence identity threshold. Differences in amino acid frequencies between lowly and highly expressed genes within the cluster are shown.

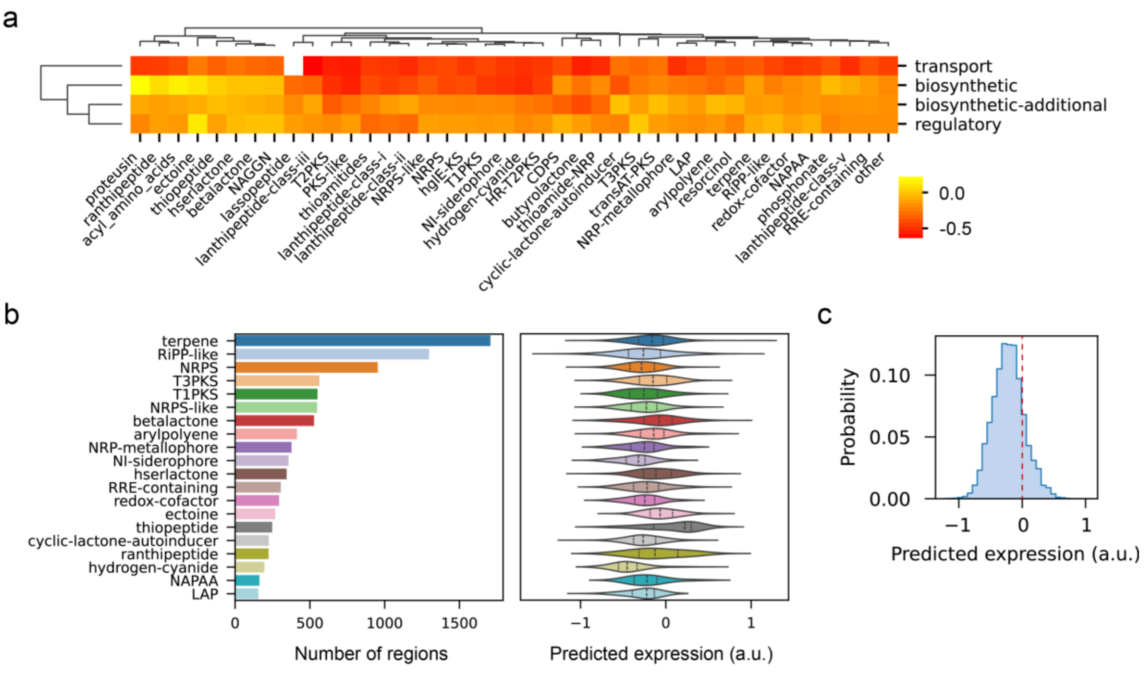


**Supplementary Figure 23.** Predicted gene expression across biosynthetic gene clusters (BGCs). **a)** Predicted gene expression levels for core biosynthetic, additional biosynthetic, regulatory, and transport genes across different BGC types. **b)** Number of regions and predicted expression levels for the top 20 BGC types. **c)** Histogram of mean expression levels of BGCs (n = 11,400).
